# Expression of amyloid-β antibody via AAV of CNS tropism alleviates Alzheimer’s disease in mice

**DOI:** 10.64898/2026.03.19.712819

**Authors:** Zhong-Min Dai, Min Jiang, Wenqing Yin, Zhounan Wang, Xiao-Jing Zhu, Mengsheng Qiu

**Affiliations:** Key Laboratory of Organ Development and Regeneration of Zhejiang Province, College of Life and Environmental Sciences, Hangzhou Normal University, Hangzhou 311121, P.R. China

**Keywords:** Amyloid-β antibody, AAV, Alzheimer’s disease

## Abstract

Alzheimer’s disease (AD), the leading cause of dementia, affects over 33 million people worldwide, with pathogenesis tied to amyloid-β (Aβ) accumulation. Although anti-Aβ monoclonal antibodies have shown clinical benefits, they often cause side effects including amyloid-related imaging abnormalities and brain microhemorrhage, especially in APOE E4 allele carriers. Here we used PHP.eB serotype adeno-associated virus (AAV), a vector with enhanced central nervous system (CNS) tropism, to deliver an Aβ antibody expression vector (AAV-LEC) into the CNS of APP/PS1 and 5×FAD mice intravenously. The AAV-LEC-mediated expression of anti-Aβ antibodies in the CNS significantly reduced the number and size of Aβ plaques at various stages in both APP/PS1 and 5×FAD mice, alongside improved spatial learning and memory. It also reversed abnormal glial activation with reduced disease-associated microglia and astrocytes, and restored oligodendrocyte differentiation and myelin formation. No brain microhemorrhage or liver damage was detected following the AAV-antibody treatment. Thus, this AAV-mediated strategy offers a promising, convenient and safe AD therapeutic approach in the future.

## INTRODUCTION

According to World Health Organization, currently more than 33 million are estimated to be affected by Alzheimer’s disease (AD), the leading cause of dementia (*1*). The amyloid cascade hypothesis holds that AD is triggered by accumulation of the amyloid-β (Aβ) peptide, the major component of amyloid plaques (*2*), followed by tau phosphorylation, neuronal cell death and eventually cognitive impairment (*3-5*). The amyloid cascade hypothesis is strongly supported by genetic studies: as autosomal dominant mutations in amyloid precursor protein (APP), presenilin 1 (PSEN1) and PSEN2 proteins that elevate the production of Aβ can cause familial AD with almost 100% penetrance (*5-9*); People who have extra copies of APP such as trisomy 21 frequently develop early onset AD (*10-12*); Various transgenic mice expressing mutated human APP or PSEN1 dominant mutants exhibit AD-like pathology (*13-15*); And the apolipoprotein E4 allele (APOE4), the strongest genetic risk factor for late onset AD, impairs Aβ clearance and accelerates Aβ aggregation (*16*); Conversely, mutations that result in decreased Aβ production can reduce the risk of AD development (*5, 17*). These genetic findings strongly support the causal role of Aβ accumulation in AD pathogenesis.

The amyloid cascade hypothesis shed the light that the progression of AD may cease if the production of Aβ is inhibited or the clearance of Aβ is enhanced (*18-20*). Over the past decades, therapeutic strategies have focused on either inhibiting Aβ production or enhancing Aβ clearance. Doubts have been raised after more than a dozen of clinical trials aiming to reduce Aβ have repetitively failed due to the lack of efficacy and toxic side effects (*21, 22*). However, Aducanumab, Lecanemab and Donanemab, three anti-Aβ monoclonal antibodies, showed clinical benefits and were approved by US Food and Drug Administration in 2021, 2023 and 2024, respectively (*23-25*). Due to the low permeability of blood brain barrier, multiple high dose anti-Aβ antibody treatments (about 10 mg/kg body weight) are required for effectively reducing Aβ plaques (*19, 25, 26*). While accumulating evidences demonstrated that anti-Aβ therapies are promising in ameliorating or even preventing the progression of AD, they all exhibited amyloid-related imaging abnormalities (ARIA) with edema or hemorrhage being the most common adverse event (*18, 23-25, 27*). Patients carrying the APOE E4 allele, especially those who are homozygous for APOE4, have a significantly increased risk of developing ARIA (*25, 28*). Effective therapies with minimal side effects are urgently needed for safe AD therapeutics.

In this study, we took advantage of the recently developed adeno-associated viruses (AAV) with enhanced CNS delivery (*29*) and the long-term durability of AAV mediated gene expression in the CNS based on a 7-year follow-up study (*30*), and utilized it to successfully deliver an Aβ antibody into the brain of AD mouse model. We demonstrated that the engineered Aβ antibody was primarily introduced into cortical neurons and its expression dramatically reduced the formation of Aβ plaques in both sexes of APP/PS1 Alzheimer’s disease mouse model. The substantial decline of Aβ plaque formation was also observed in 5×FAD mice, another AD mouse model. Moreover, the expression of Aβ antibody resulted in altered activation of microglia and astrocytes, and most importantly, the significant improvement of cognitive behavior of APP/PS1 mice.

## MATERIAL AND METHODS

### Animals

B6;C3-Tg (APPswe,PSEN1dE9)85Dbo/Mmjax (APP/PS1) transgenic mice were procured from The Jackson Laboratory and propagated through breeding with C57BL/6 mice (MMRRC Strain #034829-JAX)(*13*). Additionally, B6SJL-Tg (APPSwFlLon, PSEN1*M146L*L286V)6799Vas/Mmjax (5xFAD) mice were procured from The Jackson Laboratory and propagated through breeding with C57BL/6 mice (MMRRC Strain #034840-JAX)(*14*). The animals were accommodated in a temperature-controlled environment (22 ± 1°C) with a 12-hour light/dark cycle, where they had unrestricted access to food and water. The overall health status of the mice involved in the study was monitored through daily observations. The mice were euthanized under deep anesthesia induced by Avertin. All animal protocols adhered to ethical guidelines and were authorized by the Laboratory Animal Center and the Animal Ethics Committee of Hangzhou Normal University, China.

### Antibodies and reagents

Information for antibodies used is listed below: anti-GFAP (Oasis Biofarm, OB-PRB005), anti-IBA1 (Oasis Biofarm, OB-PRB029), anti-FLAG, anti-EGFP (Oasis Biofarm, OB-PGP003), anti-MYRF (Oasis Biofarm, OB-PGP041), anti-CC1 (Oasis Biofarm, OB-PRB070), Anti-Aβ (Cell Signaling, 15126S, IF1:200), Anti-Aβ 40/42 (Oasis Biofarm, OB-MMS049),

### Immunofluorescence Staining and analysis of images

Animals were subjected to transcardial perfusion with 4% cold paraformaldehyde (PFA) following deep anesthesia. Subsequently, the tissues were excised and underwent overnight postfixation. Cryoprotection was achieved through immersion in a 30% sucrose solution, after which the tissues were embedded in a frozen section medium (catalogue number 6502; Thermo Scientific, Waltham, MA, USA) and sectioned to a thickness of 14 μm. The immunofluorescence experimental protocol has been previously detailed. In summary, sections were initially washed with phosphate-buffered saline (PBS) and subjected to antigen retrieval in a citrate buffer at temperatures ranging from 85-98□ for 25 minutes. Following this, the sections were rinsed thrice with PBS, incubated with a blocking solution containing 10% goat serum and 0.1% Triton-X-100 for one hour, and immediately proceed to an overnight incubation at 4□ with the primary antibodies in the blocking solution. Afterward, the sections were washed three times with PBS, incubated with secondary antibodies for 2 hours at room temperature, and subsequently washed and mounted using Mowiol mounting medium with 4′,6-diamidino-2-phenylindole (DAPI) for fluorescence visualization. Fluorescent images were captured using either a ZEISS or Leica epifluorescence microscope.

### Thioflavin S staining

Thioflavin S (ThioS) staining was conducted using a 0.0025% solution of thioflavin S (Sigma, T1892-25G) in 50% ethanol for a duration of 15 minutes at room temperature. Following the staining, sections were washed sequentially in 80% and 70% ethanol for one minute each, followed by PBS, and then mounted onto slides for imaging.

### Prussian blue staining

Perl’s Prussian blue staining was employed to visualize hemosiderin deposits. Brain sections encompassing the cortex and hippocampus were analyzed. The staining procedure was carried out according to the manufacturer’s instructions (Meryer, M52355), with sections mounted on silane-coated slide glasses (Epredia Superfrost Plus Microscope Slides, 22-042-942). Imaging was performed using a slide scanner (Leica, DM6B), and subsequent analysis was conducted with the ImageJ program.

### Construction and production of pAAV vectors

To construction pAAV-LEC that expressing the Lecanemab with fused light and heavy chain, we employed de novo synthesis polymerase chain reaction (PCR). The amino acid and DNA sequence of the fused AAV-LEC is showed in supplemental information. The pAAV-LEC was then used for PHP.eB virus packaging.

### Intranasal delivery and intravenous injection

AAV viruses with PHP.eB serotype AAV-CMV-EGFP, AAV-EGFP, and AAV-LEC ordered or packaged from Shanghai Taitool Bioscience. AAV-CMV-EGFP were used to compare the CNS transduction efficiency of intranasal and intravenous administration. Wild-type mice, at the age of two months, were subjected to intranasal delivery and intravenous injection of the AAV-CMV-EGFP virus. For intranasal administration (n=3), male mice were anesthetized and positioned supine, followed by the application of 3 μl droplets to the nasal cavity using a micropipette, alternating between the right and left nostrils at one-minute intervals, amounting to a total volume of 40 μl, and a full dose of 4 × 10^11^ virus genomes (vg). Intravenous injection (n=3) was conducted via tail intravenous injection of 40 μl virus containing 4 × 10^11^ vg. APP/PS1 and 5×FAD transgenic mice received the AAV-EGFP virus as control or AAV-LEC virus for treatment by intravenous injection at a dose of 4 × 10^11^ vg.

### Aβ concentration measurement by enzyme-linked immunosorbent assay (ELISA)

Serum-free cultures of 293 cells transfected with the PAAV-LEC plasmid were pelleted at 4°C, 1,500 rpm for 10 minutes. The supernatants were collected and used as the primary antibody source. Subsequently, 100 μl of human Aβ42 (Genscript, RP10017) and human Aβ40 (Genscript, RP10004) were introduced and incubated for two hours at room temperature on a shaking device. After two hours, 200 µl of a 5% goat serum buffer was added to each well at room temperature. The positive control was Anti-Aβ 40/42 (Oasis Biofarm, OB-MMS049), while PBS served as the negative control. The cell supernatants, Anti-Aβ 40/42 (diluted in PBST–BSA), and PBS were then added and incubated for an additional two hours on a shaker. Followed by the addition of 100 μl of goat anti-human IgG (H+L) HRP-Streptavidin secondary antibody for one hour at room temperature with continuous shaking. To conclude the assay, 100 μl of 3,3′,5,5′-tetramethylbenzidine (TMB) solution (Biosharp, BL728A) was introduced as a substrate. And the reaction was stopped with 50 μl of 1 M H2SO4. Optical density (OD) values were measured at 450 nm using a Sunrise ELISA plate reader.

### Morris Water Maze (MWM) test

MWM test Spatial learning and memory evaluations were conducted within a circular tank (diameter: 100 cm; height: 40 cm) filled with opaque white water maintained at a constant temperature of 22 ± 1 °C, with a water depth of 17 cm. Mice were trained to locate a hidden platform (10 cm in diameter) submerged 1-2 cm beneath the water surface in the north-west (NW) quadrant of the tank. We employed a reference memory protocol consisting of a 5-day training regimen, during which each mouse underwent four acquisition trials daily for five consecutive days. Random starting points, excluding the target quadrant, were selected for each trial, and the mice were given 60 seconds to search for the platform. In instances where the mice failed to find the platform, they were gently directed to it and remained there for 60 seconds. The starting positions were varied daily. All trials were documented, with latency time, defined as the duration taken to find the platform, serving as an index of spatial learning. Following the training period, a memory test (probe test) was conducted by removing the platform, allowing the mice to swim freely for 60 seconds. The time spent in the target quadrant and the frequency with which the mice crossed the former platform location were recorded. The animals were then sacrificed for sampling.

### TrueGold myelin staining

Brain sections were sequentially obtained from the brains of APP/PS1 and 5XFAD mice. The tissue was fixed in 4% paraformaldehyde (PFA) and cut into 14-18 μm slices using a freezing microtome. Subsequently, the slices were brought to room temperature and baked at 37□ for 30 minutes. Myelin staining were performed using TrueGold Myelin Staining Kit (Oasis Biofarm) according to manufactures’ instruction. Briefly, tissue sections were hydrated and incubated in a 0.3% True Gold Staining solution at 45□ in the dark for 20 to 30 minutes. The sections were washed in distilled water and fixed in 1% sodium thiosulfate at 45□ for 3 minutes. After rinsing with water, dehydration through a gradient of alcohols was performed, followed by coverslipping with a mounting medium.

### Western Blot

293T cells were lysed in a buffer containing a protease inhibitor cocktail. The lysates were subsequently centrifuged at 12,000× g and 4°C for 15 minutes to remove the insoluble debris. The protein concentration of the supernatant was quantified using a BCA assay (Pierce™ BCA Protein Assay Kit, product number 23227). Protein samples were separated via 10% SDS-PAGE and transferred onto an Immobilon-P Transfer Membrane (Millipore, Kenelworth, NJ, USA). The membranes were blocked with a 5% non-fat milk solution in TBST (50 mM Tris, pH 7.4, 150 mM NaCl, and 0.1% Tween 20) for 1 hour at room temperature, followed by overnight incubation at 4°C with the respective primary antibodies diluted in the blocking buffer.

### Experimental design and statistical analyses

Two-tailed t tests were used for statistical analysis within GraphPad Prism software. The number of animals used for counting was showed in each figure.

## RESULTS

### AAV-mediated noninvasive CNS transduction for local antibody expression

To develop a highly effective and noninvasive approach for antibody delivery into the CNS, we investigated the efficiency of AAV transduction to the brain via intranasal or intravenous administration. In agreement with a previous study (*29*), our results confirmed that PHP.eB serotype AAV can be efficiently transduced into the CNS via intravenous injection (Supplemental Fig. S1). In contrast, intranasal administration failed to deliver AAV effectively into the CNS except the out layer of olfactory bulb (Supplemental Fig. S1), which suggested that other CNS regions only benefit from slow diffusion of secret proteins from olfactory bulb by intranasal routes of delivery (*31, 32*). We thus explored the intravenous injection as a more efficient and widespread transduction approach in the CNS.

AAV-LEC was constructed for CNS-tropic expression of anti-Aβ antibody for therapeutic purpose (Supplemental Fig. S2A). Successful expression of the antibody was confirmed by transient transfection of HEK293T cells followed by Western blotting (Fig. S2A). ELISA was employed to verify the specificity and affinity of antibody produced from AAV-LEC. The results demonstrated that AAV-LEC expressed antibody retained the high affinity for both Aβ40 and Aβ42 (Supplemental Fig. S2B). AAV-LEC and control AAV-EGFP were packaged as PHP.eB serotype AAV viruses, and then administered to 6-month-old APP/PS1 mice via intravenous injection. After 4 weeks of viral administration, Flag immunofluorescence was detected in both the cerebral and hippocampal regions (Supplemental Fig. S2C), indicating that AAV-LEC was able to cross the blood-brain barrier and effectively generate the target antibody in the CNS.

### AAV-LEC reduced Aβ accumulation and ameliorated cognitive impairment in AD mice

Women have a higher risk and increased severity of AD (*33*). A similar phenomenon has been observed in mouse model, for example, the male and female APP/PS1 mice begin to develop Aβ deposits by six months and three months, respectively (*34*). To assess the therapeutic efficacy of AAV-mediated antibody treatment at relatively early stages, we administrated the AAV-LEC or control AAV-EGFP viruses into approximately 6-month-old males or 3-month-old females of the APP/PS1 mice via intravenous injections. Morris Water Maze tests were conducted in 7-month-old male and female APP/PS1 mice for evaluation of their cognition and learning behaviors (Fig. 1B). Considering the reported sex differences in the AD mice, results from male and female APP/PS1 mice were analyzed separately. In both sexes, thioflavin S staining of Aβ plaques in the cerebral cortex and hippocampus revealed a substantial decline in both the number and size of Aβ plaques following the AAV-LEC administration as compared to the AAV-EGFP group (Fig. 1C-F). In the Morris Water Maze test, both male and female AAV-LEC-treated APP/PS1 mice exhibited a shorter latency to reach the hidden platform compared to AAV-EGFP-treated group during the training phase (Fig. 1G, 1L). Furthermore, AAV-LEC treated mice stayed for a longer period in the target quadrant and crossed the removed platform more frequently in the probe trial (Fig. 1I-K, 1N-P), indicating a significant enhancement of their learning and memory. These findings suggested that AAV-mediated CNS expression of anti-Aβ antibody efficiently inhibited Aβ plaque accumulation, and improved the cognitive function of AD mice.

**Figure 1.**
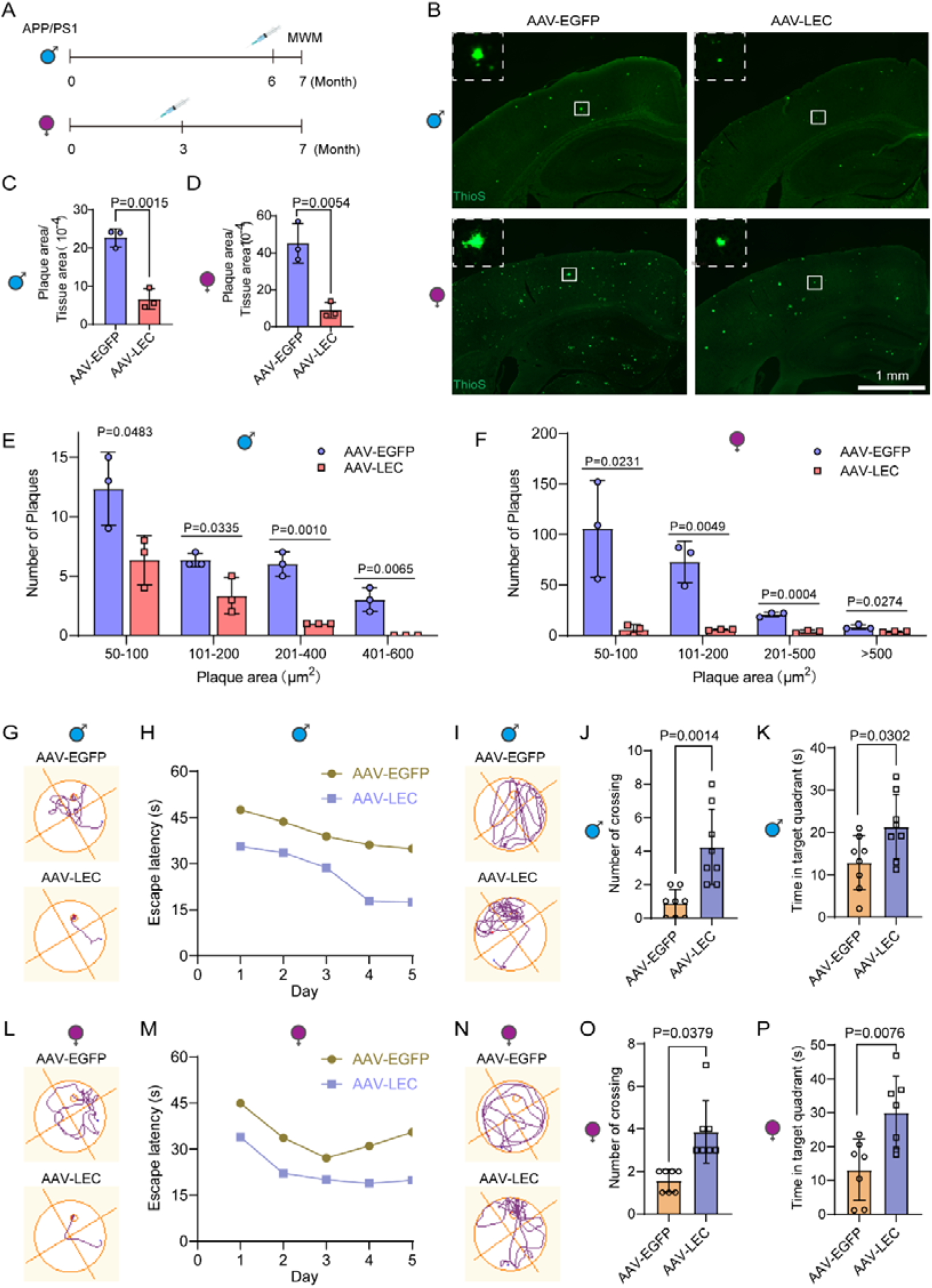
AAV-LEC reduced Aβ accumulation and ameliorated cognitive impairment in APP/PS1 mice. (**A**) Schematic AAV administration and behavioral test in male and female APP/PS1 mice. (**B**) Representative images of brain Aβ plaques stained by Thioflavin S (ThioS) in APP/PS1 male and female mice. Insets are enlarged images of dashed box region, showing the size of Aβ plaques. (**C-F**) Quantification of the Aβ plaque areas in male APP/PS1 mice (C, E) and female APP/PS1 mice (D, F). (**G-P**) Morris Water Maze (MWM) test results of male (G-K) and female (L-P) APP/PS1 mice. G, H, L and M showed the represent tracks and the time taken for the mice to locate the hidden platform during the five-day trial. I&N showed the represent tracks, and J&O showed the number of crossing the exact position where the platform was previously located. K&P showed the time spent in the quadrant where the platform was previously located.

To test if mice benefit from the therapeutic effects of AAV-LEC treatment at later stages, we administrated the AAV-LEC or control AAV-EGFP viruses into approximately 10-month-old males of the APP/PS1 mice. The MWM tests were conducted 2 months after AAV treatment. Our results showed that APP/PS1 mice treated with AAV-LEC exhibited a dramatic reduction of Aβ plaques (Fig. 2A-D). Moreover, MWM showed that AAV-LEC treatment improved the cognitive function of APP/PS1 mice (Fig. 2E-G). These results indicated that AAV-LEC treatment is still capable of improving Aβ plaque clearance and protecting neural function even after abundant plaques are formed.

**Figure 2.**
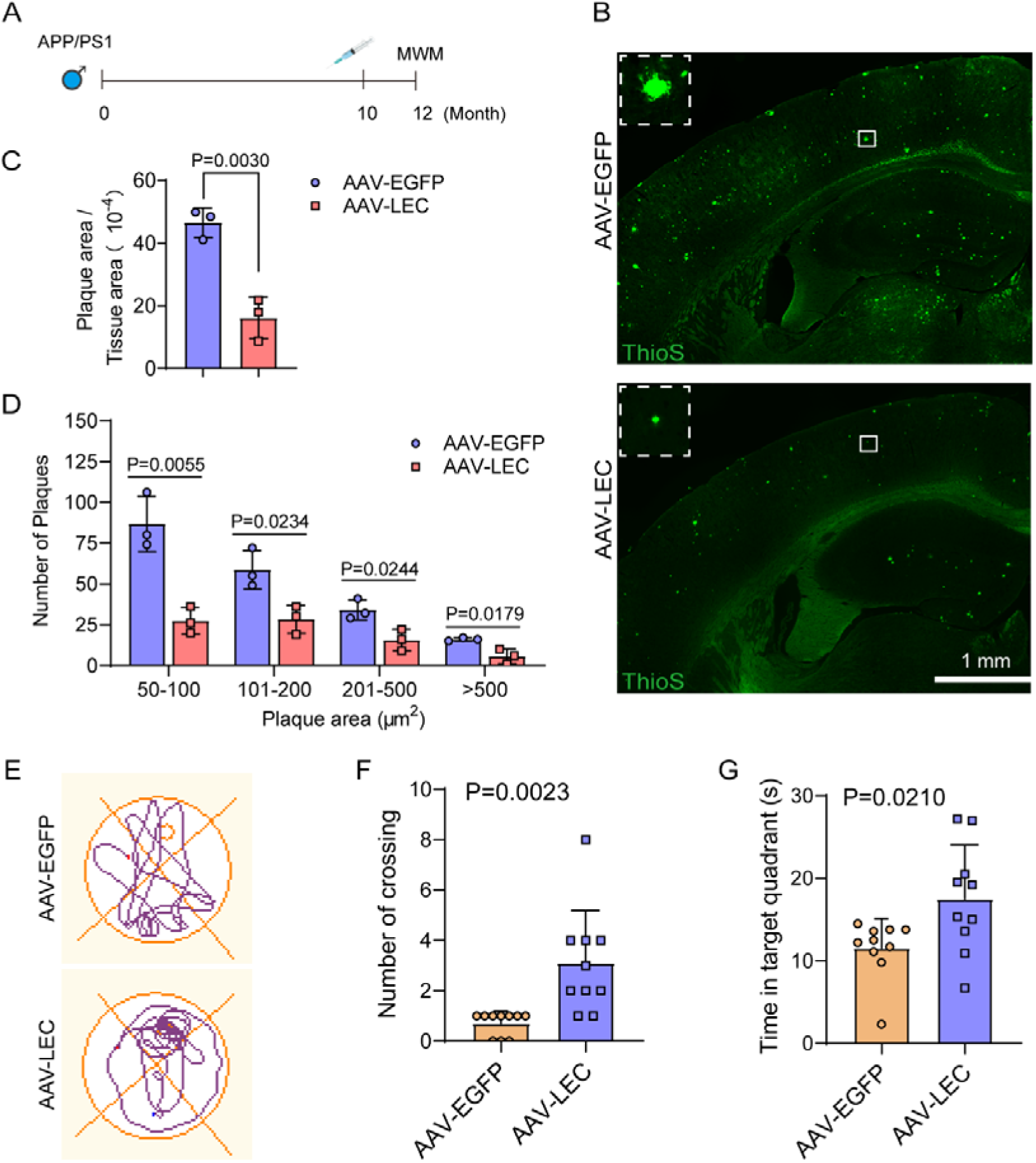
AAV-LEC cleared Aβ plaques and improved cognitive function of APP/PS1 mice. (**A**) Schematic AAV administration and behavioral test in male and female APP/PS1 mice. (**B**) Representative images of brain Aβ plaques stained by ThioS in APP/PS1 mice. Insets are enlarged images of dashed box region. (**C-D**) Quantification of the Aβ plaque areas in male APP/PS1 mice. (**E-G**) Morris water maze test showing the represent swim paths of AVV-EGFP and AAV-LEC treated APP/PS1 mice (E), the frequency of crossing the position (F) and the time spent in the quadrant (G) where the platform was previously located.

To rule out the possibility that the efficacy of AAV-LEC treatment only applies to a specific AD mouse model, we also carried out a similar study in 5×FAD AD mouse model. Following the administration of AAV-LEC virus, there was a significant decrease in both the number and the total area of Aβ plaques in the cerebral cortex and hippocampus as revealed by the ThioS staining (Fig. 3). Taken together, these findings demonstrated that AAV-LEC treatment could ameliorate the pathological symptoms in both the APP/PS1 and 5×FAD AD mouse models.

**Figure 3.**
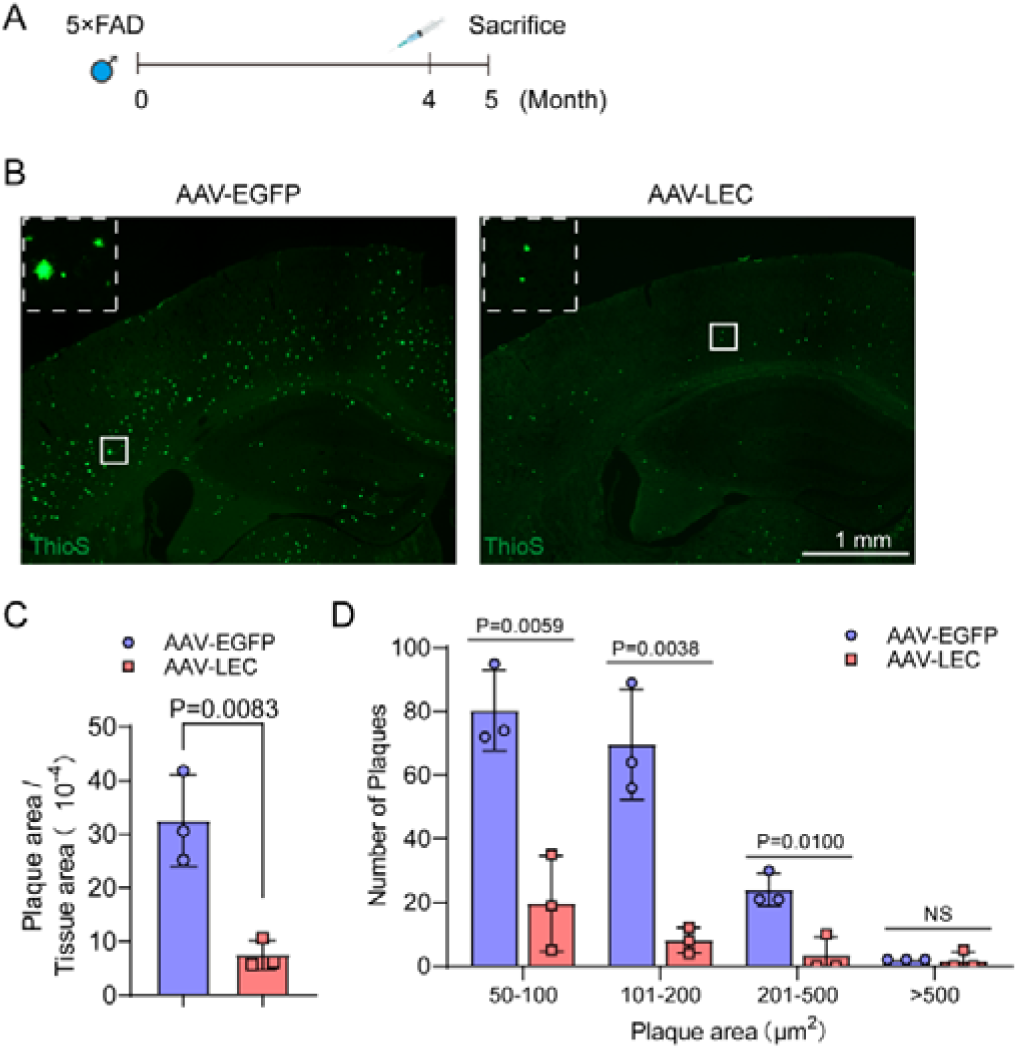
AAV-LEC reduces Aβ plaques in 5×FAD mice. (A) Schematic illustration of tail vein injection and behavioral test in male 5×FAD mice. (B) Representative images of brain Aβ plaques as stained by ThioS in 5×FAD male mice at 5 months. Insets are enlarged images of dashed box region. (C) Relative quantification of the Aβ areas in male 5×FAD mice. (D) Quantification of Aβ plaque size distribution in male 5×FAD mice.

### AAV-LEC treatment restores glial function

As previous studies showed that accumulation of Aβ in APP/PS1 mice resulted in abnormal activation of microglia and astrocytes, we therefore investigated whether AAV-LEC treatment could reduce the activation of microglia and astrocytes. By analyzing the glial cells surrounding the Aβ plaques (i.e. IBA1+ microglia and GFAP+ astrocyte), we observed a significant decline in the number of activated microglia and astrocytes following treatment with AAV-LEC in both 7- and 12-month-old APP/PS1 mice (Fig. 4 and Supplemental Fig. S3). Thus, AAV-LEC treatment led to a simultaneous decrease in Aβ accumulation and activation of astrocytes and microglia.

**Figure 4.**
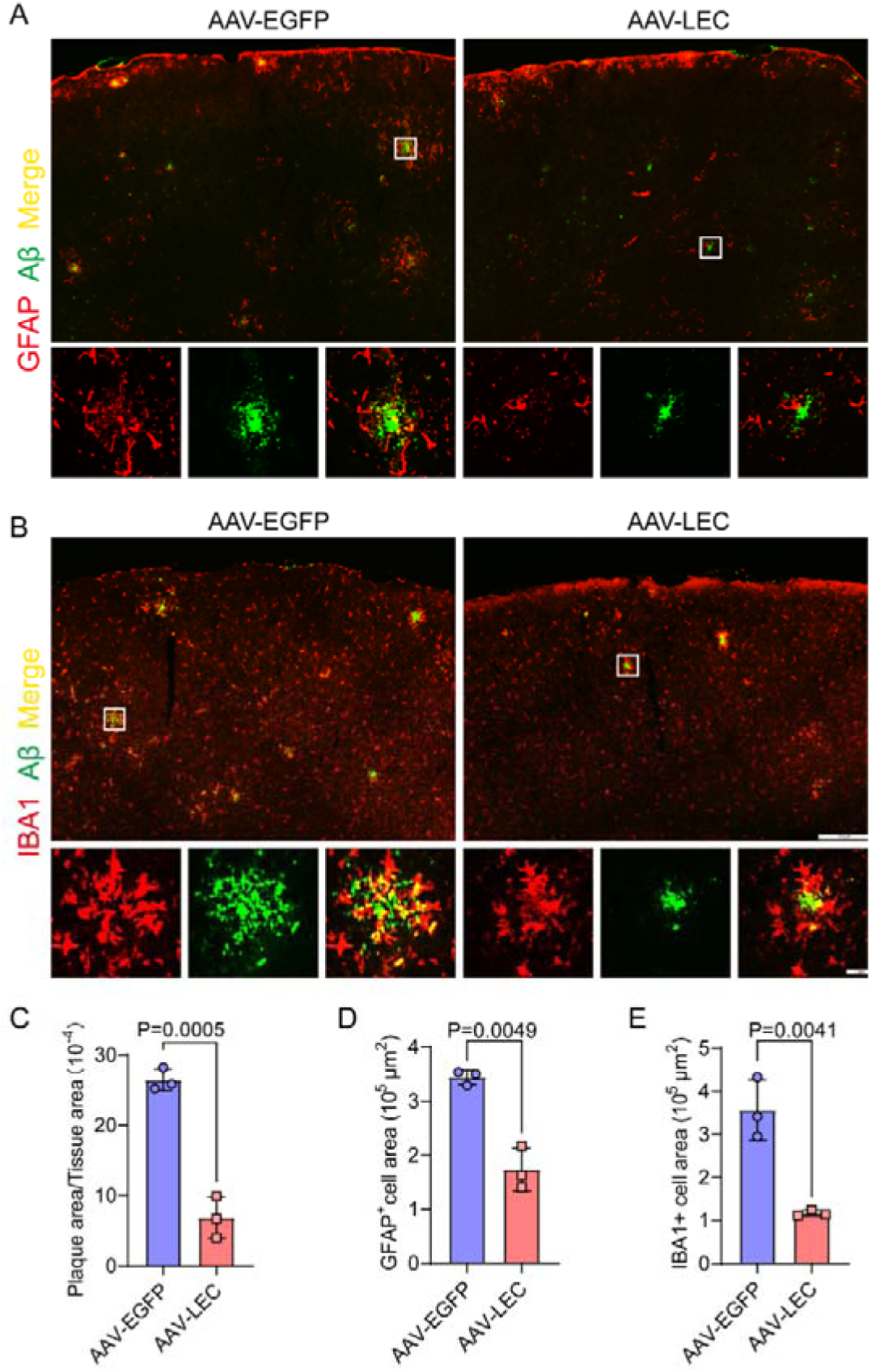
AAV-LEC treatment reduced the activation of astrocytes and microglia in APP/PS1 mice. (A) GFAP and Aβ immunofluorescence of the brain from 7-month-old AAV-EGFP and AAV-LEC treated mice. White boxes denote enlarged region. (B) IBA1 and Aβ immunofluorescence. White boxes denote enlarged region. (C) Quantification of Aβ plaque areas by anti-Aβ immunofluorescence in the cerebral cortex. (D-E) Quantification of GFAP+ astrocytes (D) and IBA1+ Microglia (E) in the cerebral cortex of APP/PS1 mice.

Since AD is associated with myelin loss, we next examined whether AAV-LEC treatment improves oligodendrocyte differentiation and myelin formation. Immunofluorescent staining demonstrated that APP/PS1 mice receiving AAV-LEC treatment elevated the number of MYRF+ and CC1+ cells to the levels of wild type mice compared to the AAV-EGFP group (Fig. 5A-C), in parallel to the increased myelinated area and myelin fiber length as revealed by TrueGold staining (Fig. 5D-F). Similar results were obtained from 12-month-old APP/PS1 mice (Supplemental Fig. S4). Collectively, these observations indicated that AAV-LEC treatment restored the differentiation of oligodendrocytes and mitigated the myelin deficit in APP/PS1 mice.

**Figure 5.**
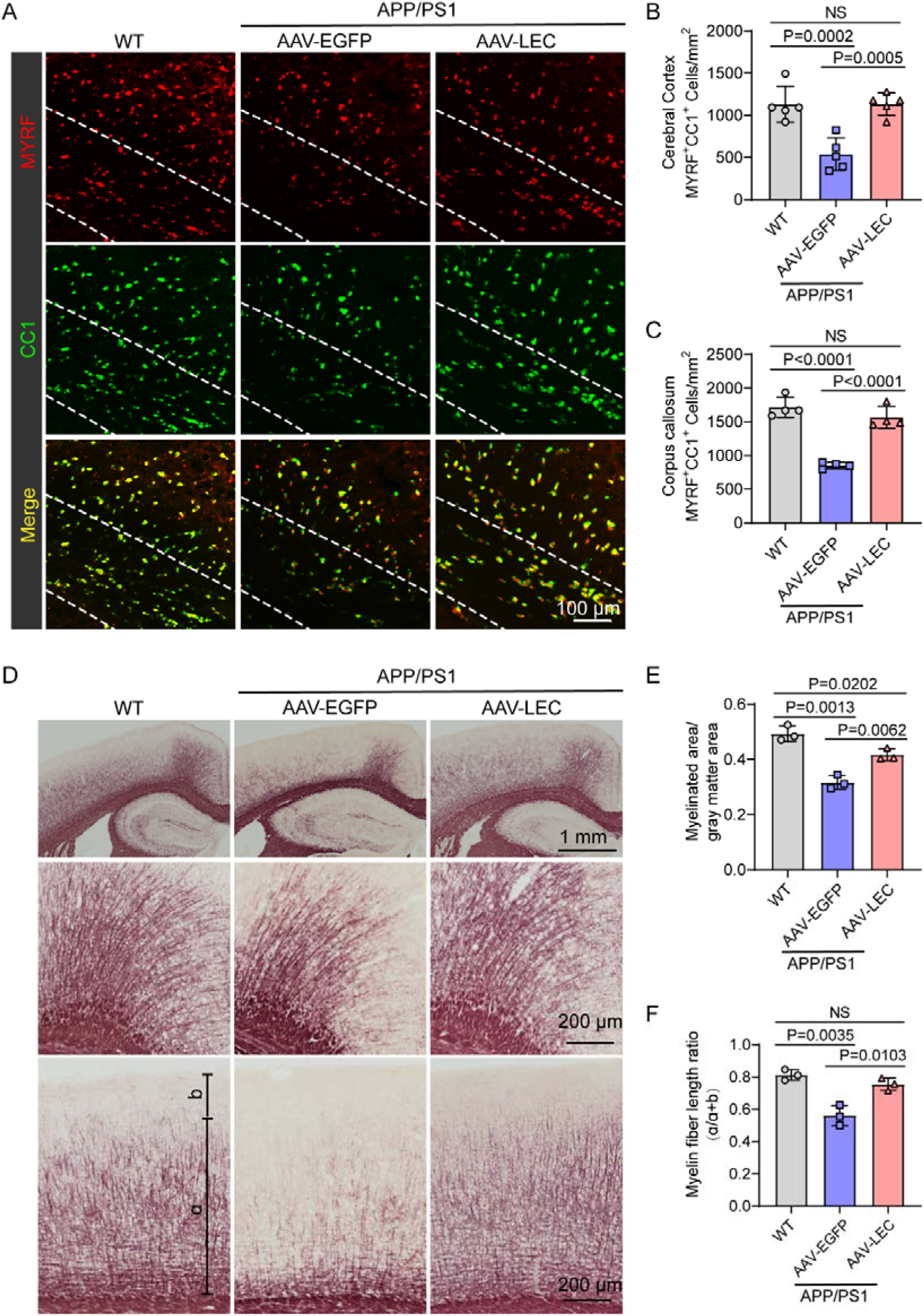
AAV-LEC treatment improves oligodendrocyte differentiation and myelin formation. (A-C) Immunofluorescence and analysis of MYRF and CC1 as differentiated oligodendrocyte marker. (D-F) Myelin staining and analysis.

### AAV-LEC treatment altered gene expression in neurodegeneration pathway

To elucidate the transcriptomic change at single cell level, the nuclei from the brain of AAV-LEC and AAV-EGFP treated APP/PS1 mice were subjected to single nucleus RNA sequencing. Based on sequence analyses, brain cells can be clustered into 7 major types by known marker genes: astrocytes (marked by *Slc1a3*), GABAergic neurons (*Gad1*), glutamatergic neurons (*Slc17a7*), microglia (*Hexb*), mature oligodendrocytes (*Aspa*), oligodendrocyte precursor cells (*Pdgfra*) and Pericytes (*Igfbp7*) (Fig. 6A-C). Compared with AAV-EGFP treatment, AAV-LEC treatment resulted in upregulation of 62 and downregulation of 84 genes (Fig. 6D, Supplemental Table 1). Six of the top 10 enriched KEGG pathways are related to neurodegeneration including AD (Fig. 6E). CellChat analyses of ligand and receptor communications between cells unveiled the diminished expression of APOE and TREM2/TYROBP after AAV-LEC treatment. Therefore, although the APP level was not significantly altered, intercellular signaling mediated by APP-TREM2/TYROBP crosstalk was greatly reduced, whereas the APP-SORL1 signaling was markedly enhanced (Supplemental Fig. S5, Fig. S6 and Supplemental Table 3). Among the four well documented AD risk genes (APP, APOE, TREM2 and SORL1) (*35, 36*), TREM2 is a microglia receptor that binds to ligands such as APP and APOE. TYROBP is the transmembrane adaptor of TREM2 involved in proliferation, phagocytosis, and inflammation (*37*). SORL1 directs trafficking of APP into recycling pathways and thus prevents the processing of APP into Aβ (*38*). The decrease of TREM2/TYROBP signaling indicated that the overall burden of Aβ is alleviated by microglial activation. Meanwhile, AAV-LEC treatment could also prevent Aβ production by promoting SORL1-mediated APP recycling.

**Figure 6.**
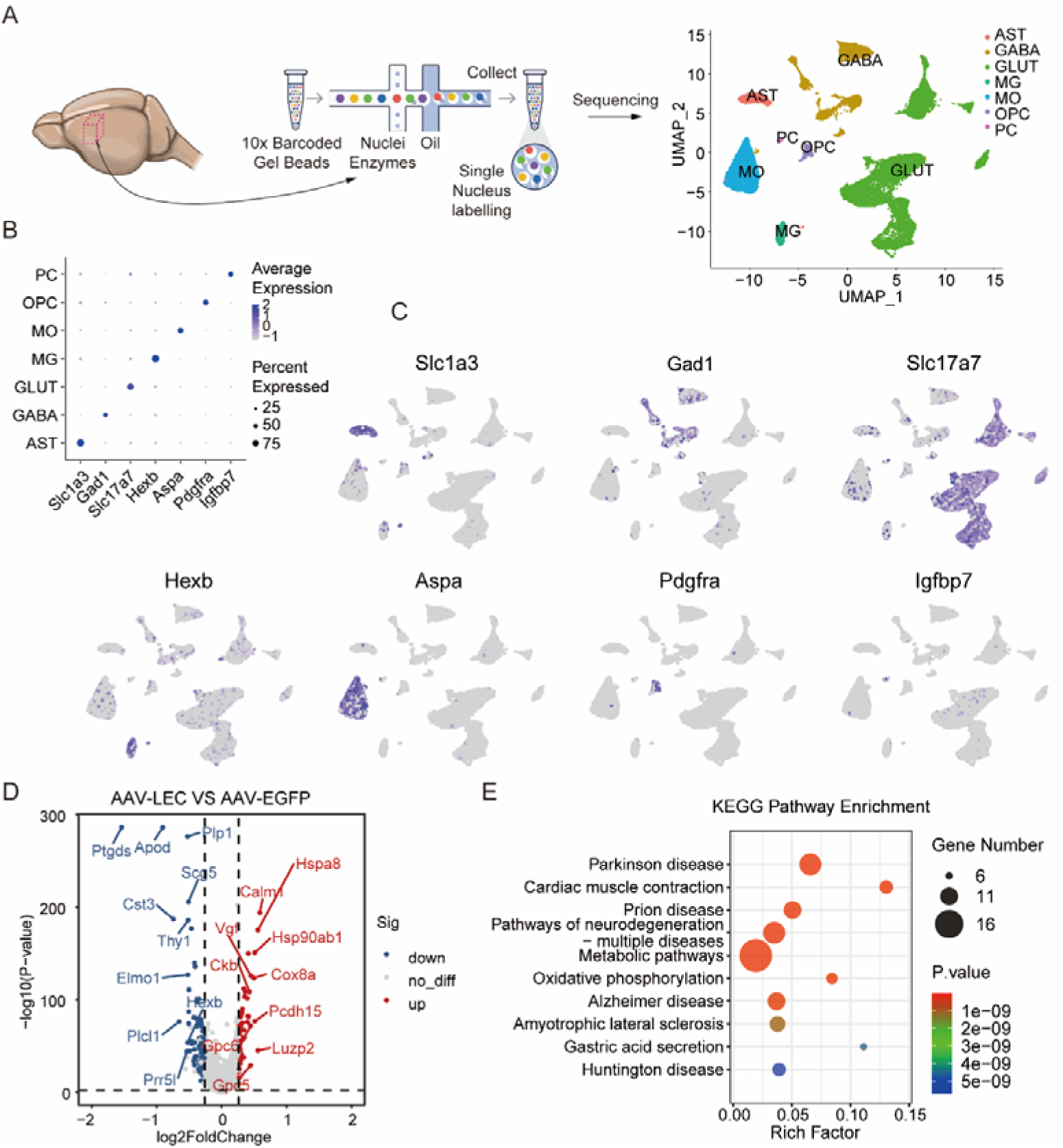
Single-nucleus sequencing revealed neurodegeneration disease related genes are enriched by AAV-LEC treatment. (A) AAV-EGFP and AAV-LEC treated APP/PS1 mice were sacrificed, the cubic regions of brain were dissected from 3 individual mice for each group and subjected to single nucleus sequencing. The cells were clustered into 7 major cell types. AST: astrocyte. GABA: GABAergic neuron. GLUT: Glutamatergic neuron. MG: microglia. MO: mature oligodendrocyte. OPC: oligodendrocyte precursor cell. PC: pericyte. (B and C) Representative marker gene expression in different cell types. (D) Volcano plot of differential expression genes from AAV-LEC treated APP/PS1 mice compared to AAV-EGFP treatment. (E) KEGG enrichment plot for the top 10 enriched pathways.

Further analysis of the differentially expressed genes showed that the major cell types except for pericytes exhibited a similar neurodegeneration-related pathway enrichment in AAV-LEC treatment group in APP/PS1 mice (Supplemental Fig. S6A-F, Supplemental Table 2 and Table 3). Notably, expression of ten genes that were previously shown to be altered in the brain of AD mice (*39-41*) was rewired back towards physiological levels. Specifically, seven genes (*Rps21*, *Hspa8*, *Ubb*, *Calm1*, *Cox6c*, *Cox8a* and *Fth1* were upregulated, and three genes (*Ptgds*, *Scg5* and *Apod*) were downregulated after AAV-LEC treatment in APP/PS1 mice (Supplemental Fig. S6G-H). These findings indicated that AAV-LEC treatment partially reversed neurodegeneration-related gene expression in APP/PS1 mice.

### AAV-LEC treatment reduces the cell proportion of disease associated glia

To gain insight into subpopulation change after AAV-LEC treatment, we further subdivided each major cell type, resulting in 6 astrocyte (AST), 18 GABAergic neuron (GABA), 17 glutamatergic neuron (GLUT), 4 microglia (MG), 8 oligodendrocyte lineage cell (OL), and 3 pericyte (PC) sub-clusters (Supplemental Table 4-9). Strikingly, the proportion of sub-clusters of microglia and astrocytes were significantly altered after AAV-LEC treatment in APP/PS1 mice (Fig. 7A-C and Fig. 8A-C).

**Figure 7.**
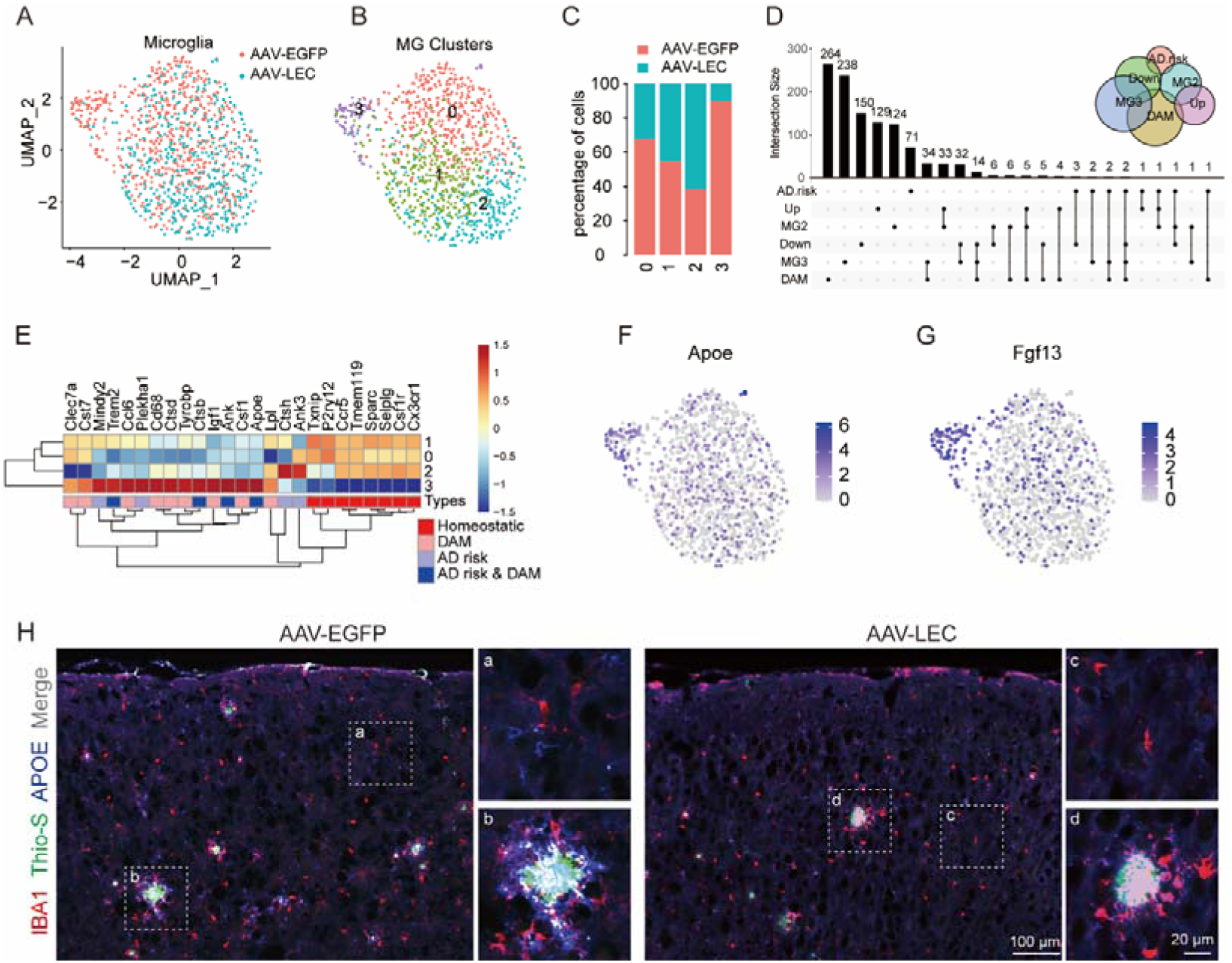
AAV-LEC treatment lead to the decrease of DAM in APP/PS1 mice. (A) Microglia from AAV-EGFP and AAV-LEC treated APP/PS1 mice. (B) Sub-clusters of microglia (MG0-MG3). (C) Fraction of cells in each sub-cluster from different treatment. AAV-LEC treatment results in increase of MG2 sub-cluster and decrease of MG3 sub-cluster. (D) Comparison of genes enriched expression in MG2, MG3, DAM, genes up- or down-regulated in microglia after AAV-LEC treatment, and AD risk genes. (E) Heatmap of homeostatic microglia and disease associated microglia (DAM) enriched genes and differentially expressed AD risk genes. (F and G) Representative expression of MG3 enriched marker gene in different MG sub-clusters. (H) APOE was significantly increased in microglia proximate to but not remote to Aβ plaque.

**Figure 8.**
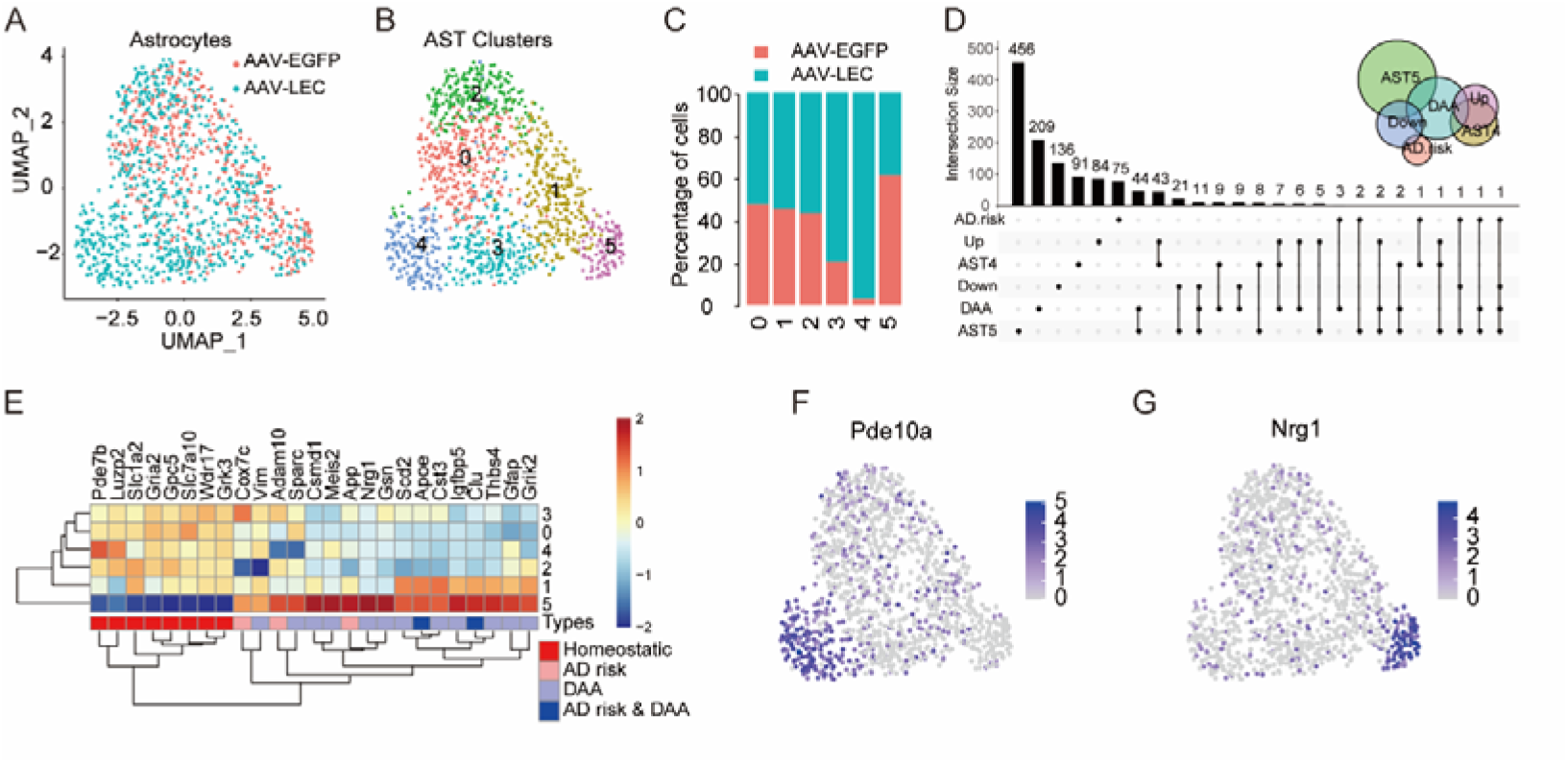
AAV-LEC treatment maintains homeostasis of astrocytes in APP/PS1 mice. (A) Astrocyte from AAV-EGFP and AAV-LEC treated APP/PS1 mice. (B) Sub-clusters of astrocytes (AST0-MG5). (C) Fraction of cells in each sub-cluster from different treatment. AAV-LEC treatment results in increase of AST4 sub-cluster and decrease of AST5 sub-cluster. (D) Comparison of genes enriched expression in AST4, AST5, DAA, genes up- or down-regulated in astrocytes after AAV-LEC treatment, and AD risk genes. (E) Heatmap of homeostatic astrocyte and disease associated astrocyte (DAA) enriched genes and differentially expressed AD risk genes. (F and G) Representative expression of AST4 (F) and AST5 (G) enriched marker gene in different AST sub-clusters.

Given that disease-associated microglia (DAM) and astrocyte (DAA) were increased in AD and aging (*42-44*), we analyzed the effects of AAV-LEC treatment on the expression of genes associated with these glia sub-clusters. It was found that the proportion of microglia sub-cluster MG3, which displayed high expression of DAM genes and AD risk genes including *Apoe*, *Trem2* and *Tyrobp*, was greatly reduced following AAV-LEC treatment (Fig. 7 A-D, Supplemental Table 7). In contrast, the proportion of MG2, which shared the characteristics of homeostatic microglia, was substantially increased (Fig. 7A-E, Supplemental Table 7). In line with the previous observation that MG3 has a higher expression level of *Apoe* (Fig. 7F), we found that APOE was more abundantly expressed in microglia proximate to Aβ plaques than in the remote ones (Fig. 7H), indicating that Aβ aggregation might directly turn homeostatic microglia into DAMs and upregulate their expression of APOE.

In addition, we found that the proportion of astrocyte sub-cluster AST5, which expressed the most enriched DAA genes and AD risk genes including *Apoe* and *Clu*, was also reduced after AAV-LEC treatment in APP/PS1 mice (Fig. 8A-E, Supplemental Table 4). APOE and CLU are ligands of TREM2 which could form complex with Aβ and serve as Aβ uptake carrier (*45*). As AST5 has the highest App expression level among all astrocyte sub-clusters, the diminished number of AST5 could be responsible for the decreased expression of App in astrocytes (Fig. 8E, Supplemental Table 1 and 4). Meanwhile, AST3 and AST4, characterized for their homeostatic gene expression pattern, were substantially increased after AAV-LEC treatment in APP/PS1 mice (Fig. 8A-E, Supplemental Table 4), suggesting that AAV-LEC treatment reversed DAA towards back to its homeostatic status.

### Safety evaluation of AAV-LEC treatment

Since ARIA are the key safety considerations in anti-Aβ immunotherapy, we used Perl’s Prussian blue staining of hemosiderin deposits to investigate whether AAV-LEC treatment increased microhemorrhage in the brain of APP/PS1 mice. Our results showed a lack of microhemorrhage in the group of AAV-LEC treated mice (Fig. 9A). Since prolonged expression of antibodies in the brain or other tissues may result in an inflammatory response, we assessed the safeties of AAV-LEC treatment by examining the liver tissue of APP/PS1 mice via HE staining and mRNA-sequencing. In comparison to the AAV-EGFP group, no significant morphological changes were observed in the liver of the AAV-LEC group (Fig. 9B). Transcriptomic analysis revealed 33 genes upregulated and 56 genes downregulated in AAV-LEC treatment (Fig. 9C and Supplemental Table 10). The most significantly enriched KEGG pathway is related to the steroid biosynthesis (Fig. 9D and Supplemental Table 11). In alignment with the reported increase of circulating cholesterol level in AD (*46*), AAV-LEC treatment significantly downregulated the expression of genes involved in steroid biosynthesis in APP/PS1 mice (Fig. 9C-F and Supplemental Table 10). The downregulation of steroid biosynthesis genes in liver was further corroborated by quantitative PCR (Fig. 9F). Taken together, our results showed that the AAV-LEC treatment did not cause microhemorrhage in the brain or damage to the liver, suggesting the safety of the AAV-mediated immunotherapy. Moreover, the reduced expression of steroid biosynthesis gene indicated that the AAV-LEC treatment improves the cholesterol homeostasis maintenance of APP/PS1 mice.

**Figure 9.**
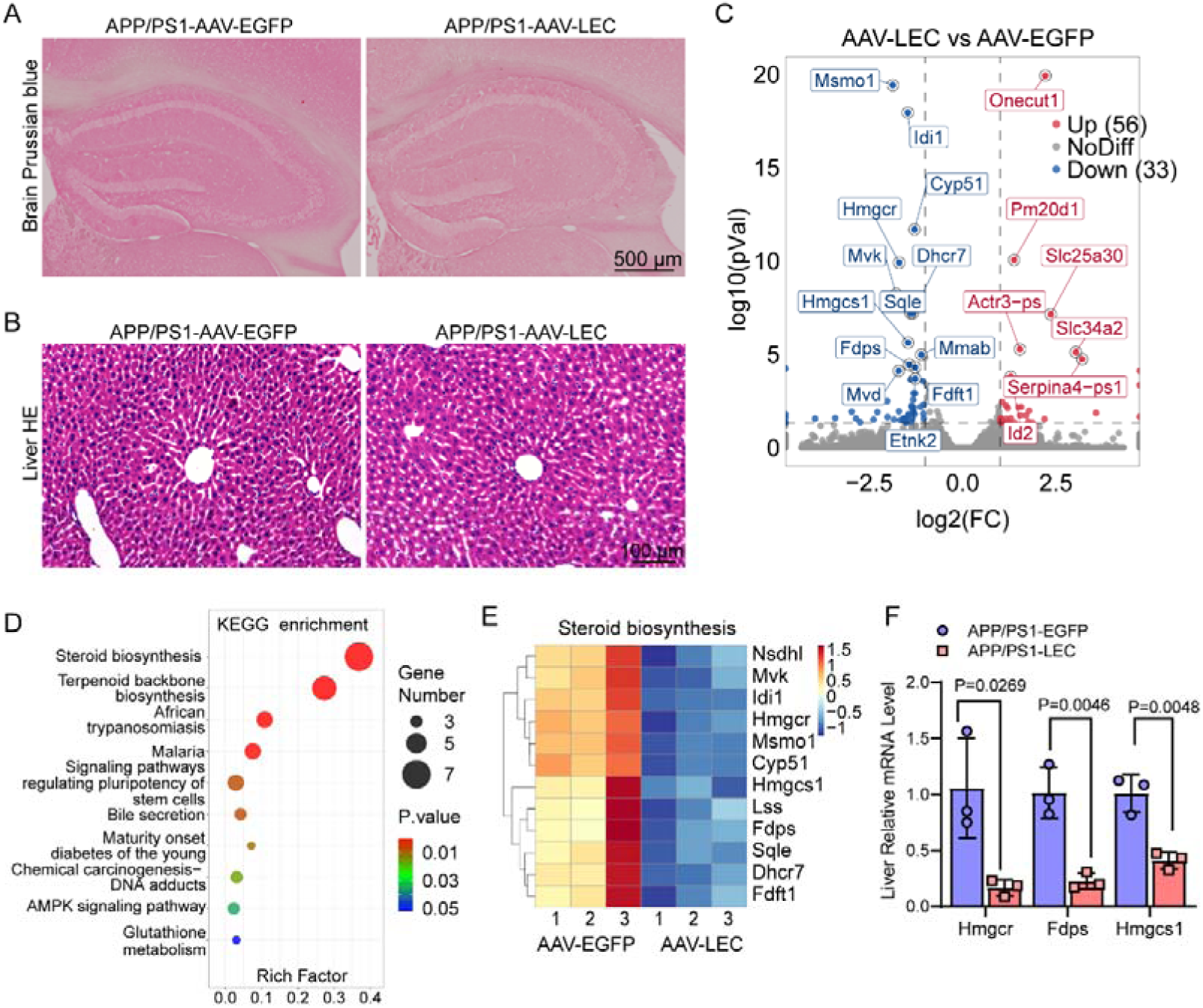
Safety evaluation of AAV-LEC treatment. (A) Perl’s Prussian blue staining of hippocampus regions in the AAV-EGFP and AAV-LEC treated 7-month old APP/PS1 mice. (B) HE staining of liver from AAV-EGFP and AAV-LEC treated APP/PS1 mice. (C) Genes significantly up and down regulated in liver RNA-seq. (D) KEGG pathway enrichment analysis of the top 10 enrichments in liver RNA-seq. (E) Heatmap analysis of 12 steroid biosynthesis genes in liver RNA-seq. (F and G) Quantitative PCR results of lipid metabolism-related genes in liver and brain tissues of APP/PS1 mice with AAV-EGFP and AAV-LEC treatment.

## DISCUSSION

In this study, we harnessed the CNS-tropism of AAV to deliver the Aβ antibody into the brain of AD mouse models, and demonstrated that systematic administration of AAV-LEC construct greatly reduced the Aβ plaques in brain and the number of disease-associated glia such as DAM and DAA, and improved the cognitive functions of AD model mice. Importantly, we did not detect microhemorrhage in the brain or damage to the liver after AAV-LEC treatment, suggesting that AAV-mediated CNS delivery of vectors expressing therapeutic antibodies could be a promising and safe immunotherapy for AD and other protein aggregation-associated neurodegeneration diseases.

Recent studies demonstrated that the proportions of DAM and DAA arise and increase along AD progression (*42-44*). Our results revealed that APOE, TREM2 and SPP1 genes were enriched in MG3 sub-clusters (Fig. 7E and Supplemental Table 7), whereas APOE, CLU, APP and AMAM10 were enriched in AST5 sub-clusters (Fig. 8E and Supplemental Table 4). It was previously shown that APOE can enhance APP transcription in neurons, leading to increased Aβ production and accumulation (*16*), while APOE and CLU together promote Aβ uptake and clearance via TREM2 (*45*). ADAM10 acts as α secretase that processes APP in a non-Aβ way (*47*). SPP1 was recently identified as a factor induced by Lecanemab treatment that promotes Aβ clearance (*48*). Therefore, DAM and DAA appear to possess dual roles in AD progression, and could be both beneficial and detrimental to the survival of neurons. However, the reduced proportions of MG3 and AST5 sub-clusters observed in APP/PS1 mice after AAV-LEC treatment might be secondary to the decreased Aβ burden, as the antibody mediated Aβ clearance exceeds Aβ accumulation.

AAV is a replication-incompetent parvovirus whose propagation is helper-dependent, which stimulates relatively mild immune response compared with other viruses. AAV mediated transgene expression can last for at least 7 years in the CNS (*30*). Recently, a few engineered AAV serotypes can be used for CNS delivery via systematic administration in rodents and non-human primates (*29, 49-53*). Here we used the PHP.eB serotype AAV, a well characterized and highly efficient serotype for CNS transduction with lowered liver transduction in B6/C57 mouse line (*29*). We found that intravenous viral administration allows efficient antibody expression in the CNS (Supplemental Fig. S2), whereas intranasal administration only transduce AAV into the out layer of olfactory bulb (Supplemental Fig. S1), demonstrating that intravenous administration is much more efficient than intranasal administration. Considering that promoters that expressed transgene at physiological level or in desired cells showed better safety (*54*), we chose the EF1α promoter, a broad and moderately active promoter, to express the antibody in order to avoid intense overexpression. Accumulating evidences showed that all anti-Aβ injection therapies shared a high prevalence of ARIA as a side effect (*18, 23-25, 27*), and high dose of AAV administration in clinical trials may trigger liver toxicity. However, we haven’t detected apparent side effects observed in clinical antibody injection, such as brain microhemorrhage or liver damage in our study. Another advantage with the AAV approach is the long-lasting effect of one-time treatment, instead of multiple regular injections used in the current antibody therapy. Together, our findings strongly suggest that the CNS tropism AAV-mediated moderate expression of antibody may represent a safer, easier and possibly more effective alternative approach for the therapeutic treatment of AD in the future.

## ACKNOWLEDGMENTS

This work was supported by the Ministry of Science and Technology China Brain Initiative Grant (Grant No. 2022ZD0204701), the National Natural Science Foundation of China (Grant No. 31871480 and 32371022) and Zhejiang Provincial Natural Science Foundation of China (Grant No. LY24C090007).

## AUTHOR CONTRIBUTIONS

Z.M.D. and M.Q. designed the research plan. Z.M.D., W.G., D.Y., X.J.Z., S.S., H.H., M.J. and B.X. performed all the experiments and analyzed the data. Z.Z. provided resources and inputs. Z.M.D. and Q.M. wrote the manuscript. Q.M. supervised the project.

## REFERENCES

1. WHO. (2025).

2. G. G. Glenner, C. W. Wong, Alzheimer’s disease: initial report of the purification and characterization of a novel cerebrovascular amyloid protein. Biochem Biophys Res Commun 120, 885–890 (1984).

3. J. A. Hardy, G. A. Higgins, Alzheimer’s disease: the amyloid cascade hypothesis. Science 256, 184–185 (1992).

4. G. B. Frisoni et al., The probabilistic model of Alzheimer disease: the amyloid hypothesis revised. Nat Rev Neurosci 23, 53–66 (2022).

5. R. Sims, M. Hill, J. Williams, The multiplex model of the genetics of Alzheimer’s disease. Nat Neurosci 23, 311–322 (2020).

6. M. C. Chartier-Harlin et al., Early-onset Alzheimer’s disease caused by mutations at codon 717 of the beta-amyloid precursor protein gene. Nature 353, 844–846 (1991).

7. A. Goate et al., Segregation of a missense mutation in the amyloid precursor protein gene with familial Alzheimer’s disease. Nature 349, 704–706 (1991).

8. J. Murrell, M. Farlow, B. Ghetti, M. D. Benson, A mutation in the amyloid precursor protein associated with hereditary Alzheimer’s disease. Science 254, 97–99 (1991).

9. C. Reitz, M. A. Pericak-Vance, T. Foroud, R. Mayeux, A global view of the genetic basis of Alzheimer disease. Nat Rev Neurol 19, 261–277 (2023).

10. F. K. Wiseman et al., A genetic cause of Alzheimer disease: mechanistic insights from Down syndrome. Nat Rev Neurosci 16, 564–574 (2015).

11. P. Mumford et al., Genetic Mapping of APP and Amyloid-beta Biology Modulation by Trisomy 21. J Neurosci 42, 6453–6468 (2022).

12. A. Rovelet-Lecrux et al., APP locus duplication causes autosomal dominant early-onset Alzheimer disease with cerebral amyloid angiopathy. Nat Genet 38, 24–26 (2006).

13. J. L. Jankowsky et al., Mutant presenilins specifically elevate the levels of the 42 residue beta-amyloid peptide in vivo: evidence for augmentation of a 42-specific gamma secretase. Hum Mol Genet 13, 159–170 (2004).

14. H. Oakley et al., Intraneuronal beta-amyloid aggregates, neurodegeneration, and neuron loss in transgenic mice with five familial Alzheimer’s disease mutations: potential factors in amyloid plaque formation. J Neurosci 26, 10129–10140 (2006).

15. H. Sasaguri et al., APP mouse models for Alzheimer’s disease preclinical studies. EMBO J 36, 2473–2487 (2017).

16. Y. A. Huang, B. Zhou, M. Wernig, T. C. Sudhof, ApoE2, ApoE3, and ApoE4 Differentially Stimulate APP Transcription and Abeta Secretion. Cell 168, 427-441 e421 (2017).

17. T. Jonsson et al., A mutation in APP protects against Alzheimer’s disease and age-related cognitive decline. Nature 488, 96–99 (2012).

18. X. Gu et al., Monoclonal antibody therapy for Alzheimer’s disease focusing on intracerebral targets. Biosci Trends, (2024).

19. M. Neatu et al., Monoclonal Antibody Therapy in Alzheimer’s Disease. Pharmaceutics 16, (2023).

20. E. Karran, B. De Strooper, The amyloid hypothesis in Alzheimer disease: new insights from new therapeutics. Nat Rev Drug Discov 21, 306–318 (2022).

21. F. Panza, M. Lozupone, G. Logroscino, B. P. Imbimbo, A critical appraisal of amyloid-beta-targeting therapies for Alzheimer disease. Nat Rev Neurol 15, 73–88 (2019).

22. Y. Zhang, H. Chen, R. Li, K. Sterling, W. Song, Amyloid beta-based therapy for Alzheimer’s disease: challenges, successes and future. Signal Transduct Target Ther 8, 248 (2023).

23. J. Sevigny et al., The antibody aducanumab reduces Abeta plaques in Alzheimer’s disease. Nature 537, 50–56 (2016).

24. C. H. van Dyck et al., Lecanemab in Early Alzheimer’s Disease. N Engl J Med 388, 9–21 (2023).

25. S. Salloway et al., TRAILBLAZER-ALZ 4: A phase 3 trial comparing donanemab with aducanumab on amyloid plaque clearance in early, symptomatic Alzheimer’s disease. Alzheimers Dement 21, e70293 (2025).

26. W. M. Pardridge, Treatment of Alzheimer’s Disease and Blood-Brain Barrier Drug Delivery. Pharmaceuticals (Basel) 13, (2020).

27. L. S. Honig et al., ARIA in patients treated with lecanemab (BAN2401) in a phase 2 study in early Alzheimer’s disease. Alzheimers Dement (N Y) 9, e12377 (2023).

28. S. Y. Jeong et al., Incidence of Amyloid-Related Imaging Abnormalities in Phase III Clinical Trials of Anti-Amyloid-beta Immunotherapy: An Updated Meta-Analysis. Neurology 104, e213483 (2025).

29. K. Y. Chan et al., Engineered AAVs for efficient noninvasive gene delivery to the central and peripheral nervous systems. Nat Neurosci 20, 1172–1179 (2017).

30. S. Marco et al., Seven-year follow-up of durability and safety of AAV CNS gene therapy for a lysosomal storage disorder in a large animal. Mol Ther Methods Clin Dev 23, 370–389 (2021).

31. L. R. Belur et al., Intranasal Adeno-Associated Virus Mediated Gene Delivery and Expression of Human Iduronidase in the Central Nervous System: A Noninvasive and Effective Approach for Prevention of Neurologic Disease in Mucopolysaccharidosis Type I. Hum Gene Ther 28, 576–587 (2017).

32. D. A. Wolf et al., Lysosomal enzyme can bypass the blood-brain barrier and reach the CNS following intranasal administration. Mol Genet Metab 106, 131–134 (2012).

33. N. T. Aggarwal, M. M. Mielke, Sex Differences in Alzheimer’s Disease. Neurol Clin 41, 343–358 (2023).

34. G. D. Manocha et al., Temporal progression of Alzheimer’s disease in brains and intestines of transgenic mice. Neurobiol Aging 81, 166–176 (2019).

35. C. Bellenguez et al., New insights into the genetic etiology of Alzheimer’s disease and related dementias. Nat Genet 54, 412–436 (2022).

36. N. Karagas, J. E. Young, E. E. Blue, S. Jayadev, The Spectrum of Genetic Risk in Alzheimer Disease. Neurol Genet 11, e200224 (2025).

37. J. V. Haure-Mirande, M. Audrain, M. E. Ehrlich, S. Gandy, Microglial TYROBP/DAP12 in Alzheimer’s disease: Transduction of physiological and pathological signals across TREM2. Mol Neurodegener 17, 55 (2022).

38. E. Rogaeva et al., The neuronal sortilin-related receptor SORL1 is genetically associated with Alzheimer disease. Nat Genet 39, 168–177 (2007).

39. R. P. Raju, L. Cai, A. Tyagi, S. Pugazhenthi, Interactions of Cellular Energetic Gene Clusters in the Alzheimer’s Mouse Brain. Mol Neurobiol 61, 476–486 (2024).

40. H. Mathys et al., Single-cell transcriptomic analysis of Alzheimer’s disease. Nature 570, 332–337 (2019).

41. E. Miyoshi et al., Spatial and single-nucleus transcriptomic analysis of genetic and sporadic forms of Alzheimer’s disease. Nat Genet 56, 2704–2717 (2024).

42. N. Habib et al., Disease-associated astrocytes in Alzheimer’s disease and aging. Nat Neurosci 23, 701–706 (2020).

43. H. Keren-Shaul et al., A Unique Microglia Type Associated with Restricting Development of Alzheimer’s Disease. Cell 169, 1276–1290 e1217 (2017).

44. L. Lin et al., Disease-associated astrocytes and microglia markers are upregulated in mice fed high fat diet. Sci Rep 13, 12919 (2023).

45. F. L. Yeh, Y. Wang, I. Tom, L. C. Gonzalez, M. Sheng, TREM2 Binds to Apolipoproteins, Including APOE and CLU/APOJ, and Thereby Facilitates Uptake of Amyloid-Beta by Microglia. Neuron 91, 328–340 (2016).

46. F. Yin, Lipid metabolism and Alzheimer’s disease: clinical evidence, mechanistic link and therapeutic promise. FEBS J 290, 1420–1453 (2023).

47. P. H. Kuhn et al., ADAM10 is the physiologically relevant, constitutive alpha-secretase of the amyloid precursor protein in primary neurons. EMBO J 29, 3020–3032 (2010).

48. G. Albertini et al., The Alzheimer’s therapeutic Lecanemab attenuates Abeta pathology by inducing an amyloid-clearing program in microglia. Nat Neurosci, (2025).

49. T. F. Shay et al., Primate-conserved carbonic anhydrase IV and murine-restricted LY6C1 enable blood-brain barrier crossing by engineered viral vectors. Sci Adv 9, eadg6618 (2023).

50. B. E. Deverman et al., Cre-dependent selection yields AAV variants for widespread gene transfer to the adult brain. Nat Biotechnol 34, 204–209 (2016).

51. D. Goertsen et al., AAV capsid variants with brain-wide transgene expression and decreased liver targeting after intravenous delivery in mouse and marmoset. Nat Neurosci 25, 106–115 (2022).

52. X. Chen et al., Engineered AAVs for non-invasive gene delivery to rodent and non-human primate nervous systems. Neuron 110, 2242–2257 e2246 (2022).

53. M. R. Chuapoco et al., Adeno-associated viral vectors for functional intravenous gene transfer throughout the non-human primate brain. Nat Nanotechnol 18, 1241–1251 (2023).

54. A. M. Keeler, W. Zhan, S. Ram, K. A. Fitzgerald, G. Gao, The curious case of AAV immunology. Mol Ther 33, 1946–1965 (2025).

