## Supplemental information for "Expression of amyloid-β antibody via AAV of CNS tropism alleviates Alzheimer’s disease in mice"

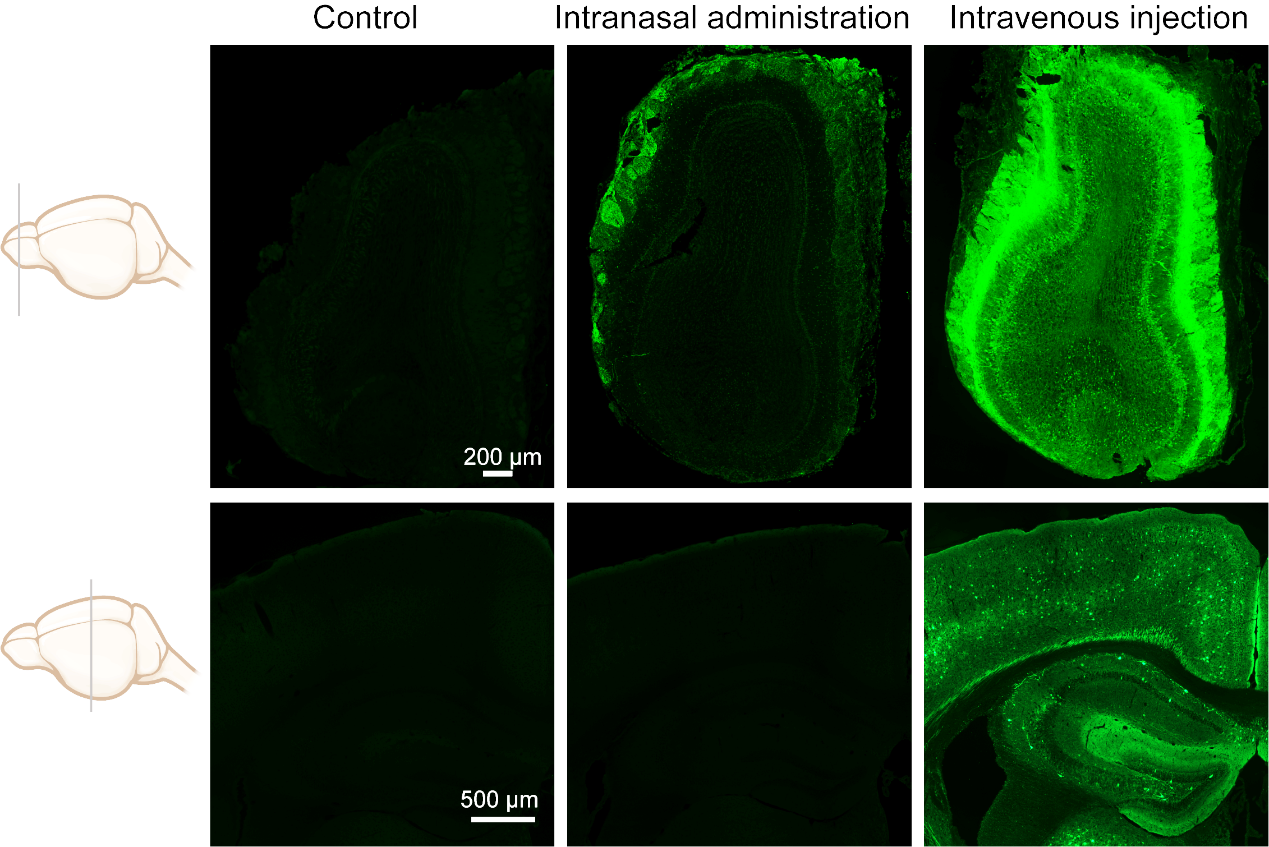

**Figure S1. Efficient transduction of AAV-PHP.eB by intravenous injection.** Sections of olfactory bulb and cerebral context were used to examine the transduction of PHP.eB serotype AAV. No EGFP signals were observed without AAV-CMV-EGFP administration (Control). Intravenous injection of AAV-CMV-EGFP resulted in highly efficient expression of EGFP in the olfactory bulb and cerebral context, whereas intranasal administration only produced EGFP expression in the outer layer olfactory bulb.

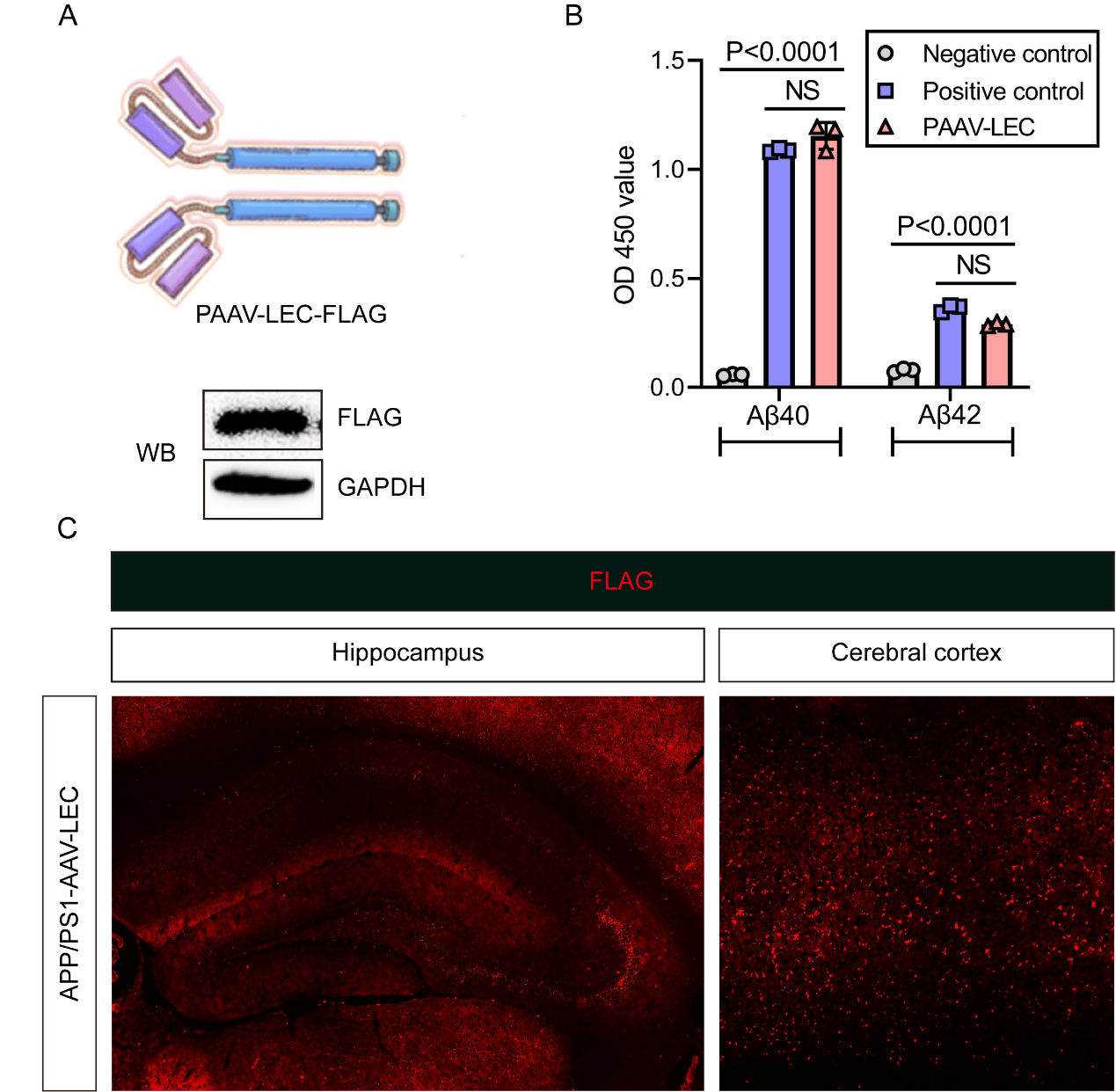

**Figure S2. Validation of AAV-LEC for CNS expression.** (A) Top, light chain variant from Lecanemab was fused to heavy chain and cloned into AAV vector. Bottom, western blotting to verify the expression of fused antibody. (B) The fused antibody retained its specificity and binding activity. Aβ40 or Aβ42 coated plate was subjected to ELISA. (C) Immunofluorescent analysis of FLAG after AAV-LEC administration.

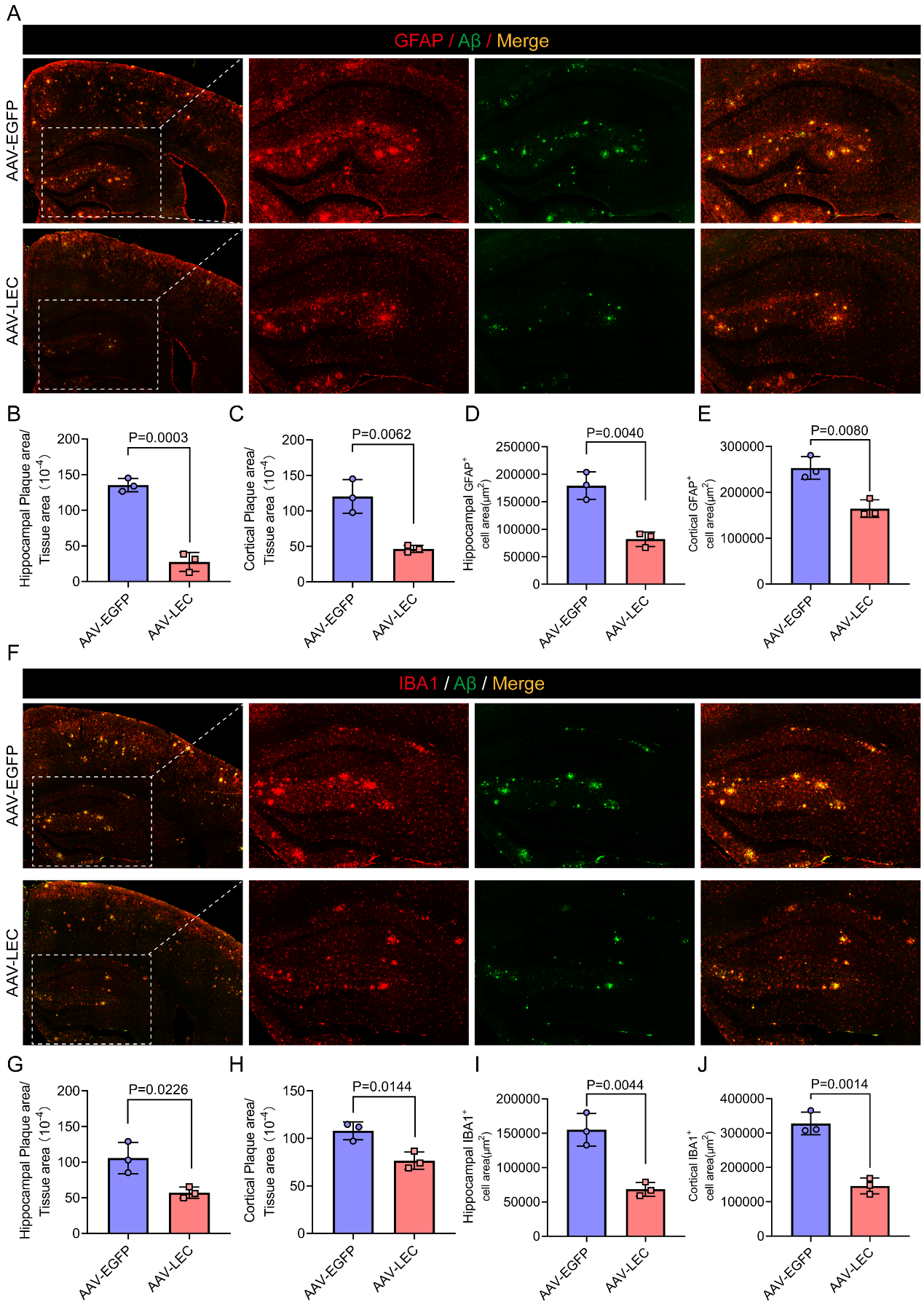

**Figure S3. AAV-LEC treatment reduced the Aβ plaques and the activation of astrocytes and microglia in APP/PS1 mice.** (A) GFAP and Aβ immunofluorescence of the brain from 12-month-old AAV-EGFP and AAV-LEC treated APP/PS1 mice. (B-E) Quantification of Aβ plaque areas and GFAP+ astrocytes. (F) IBA1 and Aβ immunofluorescence. (G-J) Quantification of Aβ plaque areas and IBA1+ microglia.

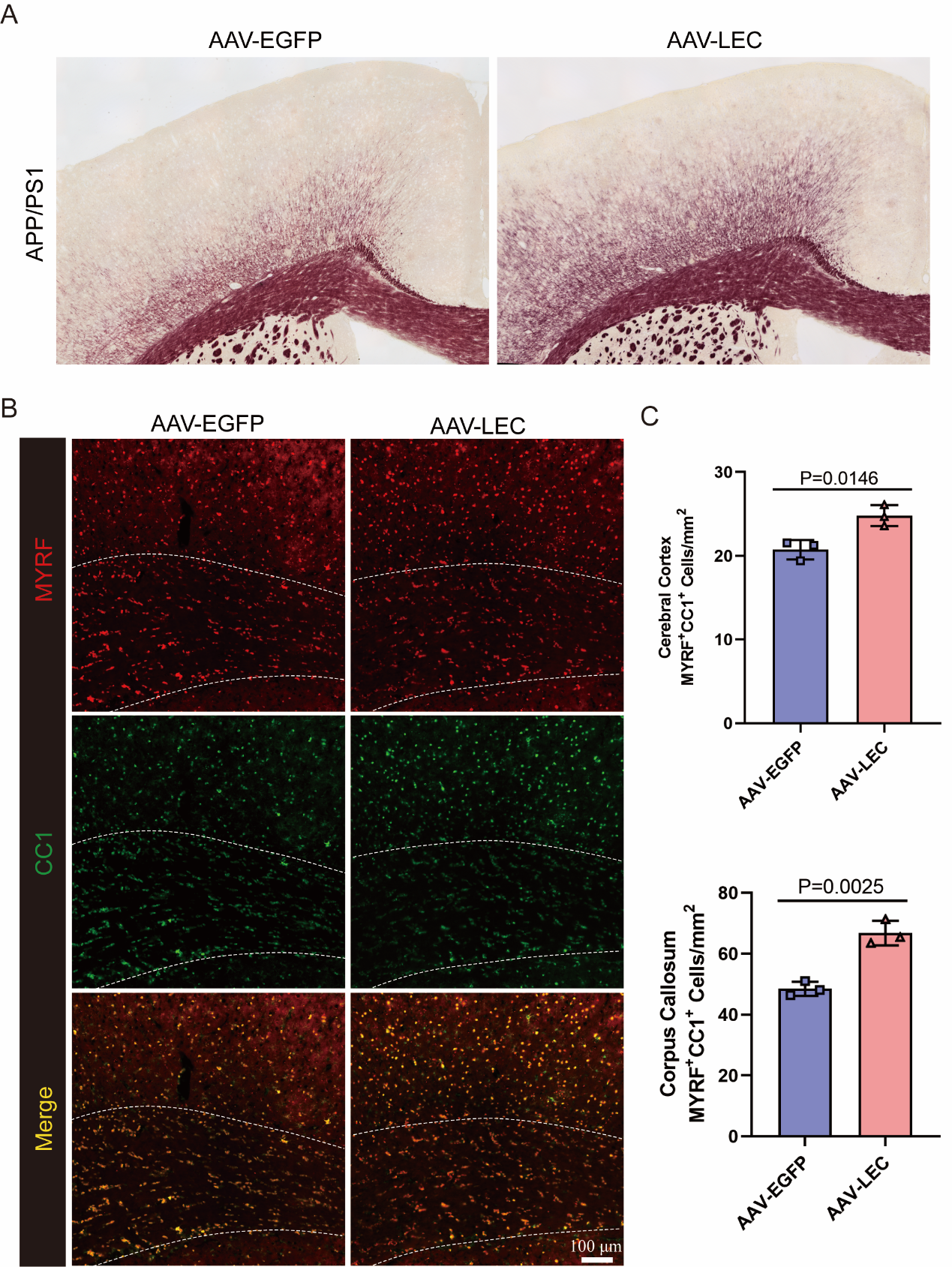

**Figure S4. AAV-LEC treatment improved myelin formation.** (A) TrueGold staining of the brain tissues from 12-month-old AAV-EGFP and AAV-LEC treated APP/PS1 mice. (B, C) Immunofluorescent staining with MYRF and CC1 antibodies, and statistical analyses of positive cells in the brain of 12-month-old APP/PS1 mice treated with AAV-EGFP and AAV-LEC.

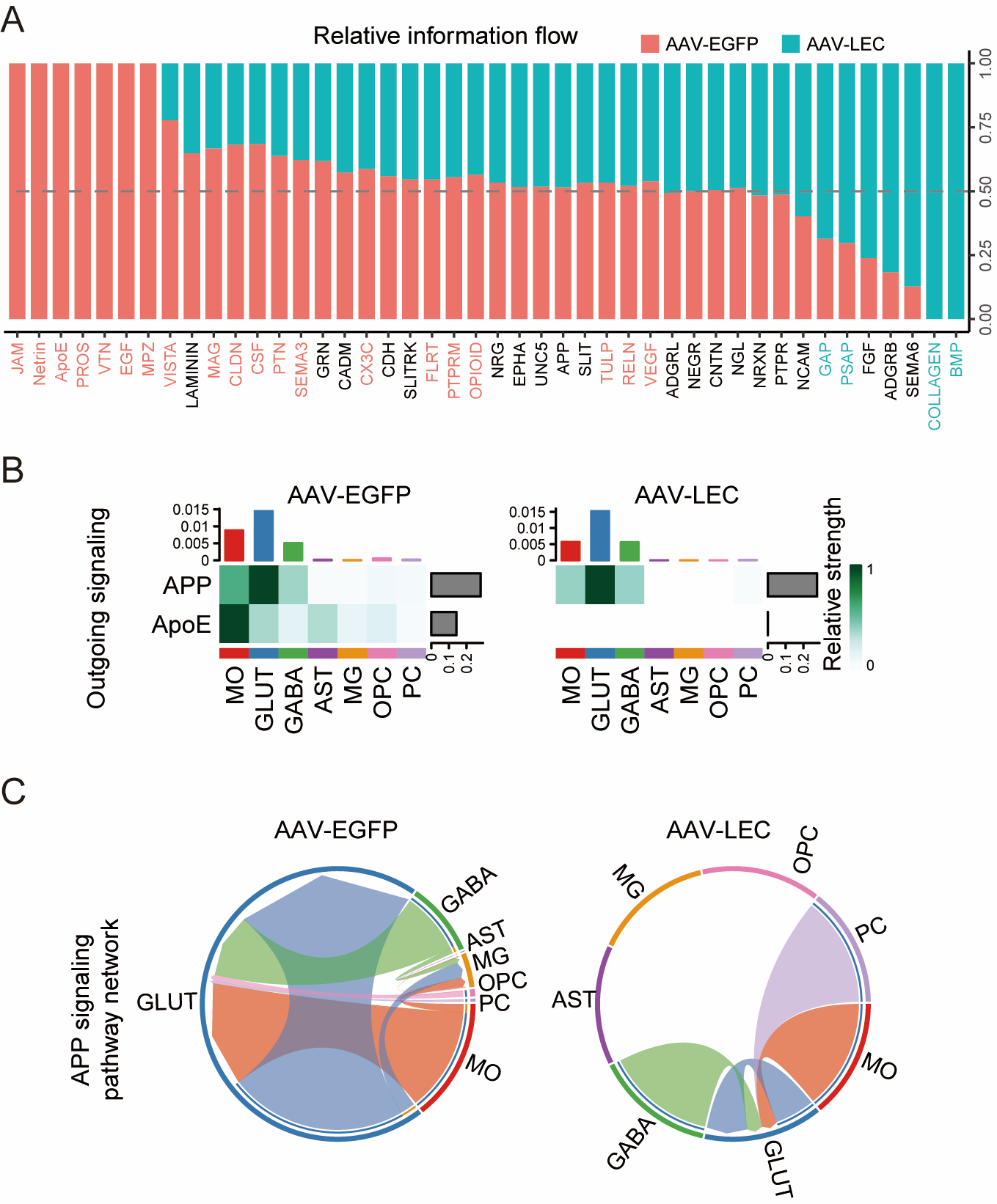

**Figure S5. AAV-LEC treatment altered intercellular crosstalk in APP/PS1 mice.** (A) AAV-LEC treatment altered information flow in different cell types from the brain of APP/PS1 mice. (B) The outgoing signaling strength of APP and APOE from various cells in AAV-EGFP and AAV-LEC treated APP/PS1 mice. (C) AAV-LEC treatment resulted in significantly altered APP signaling pathway network among various cell types.

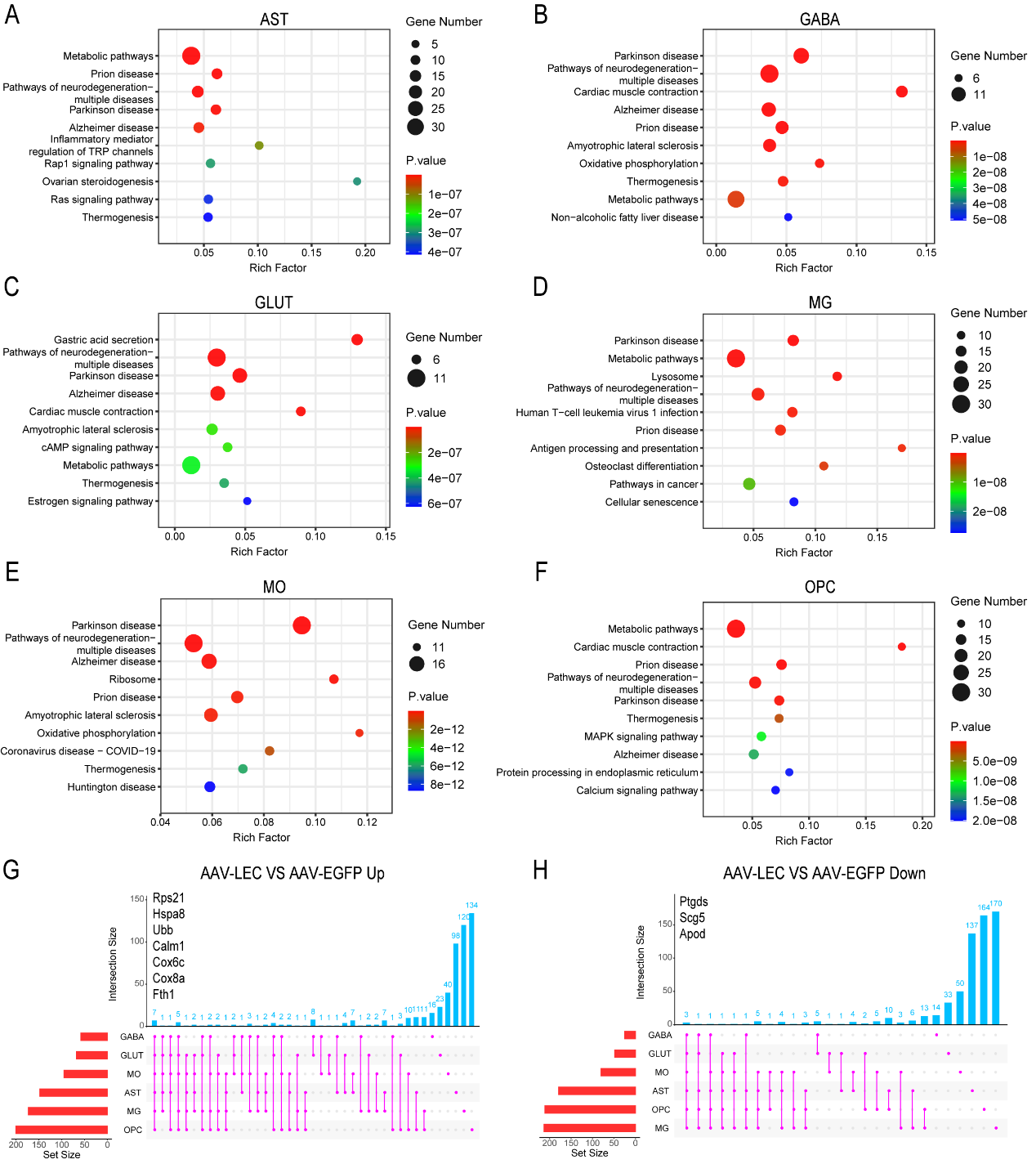

**Figure S6. Enrichment of pathways for AD and other neurodegenerative diseases in various cell types in APP/PS1 mice after AAV-LEC treatment.** (A-F) KEGG pathway enrichment plot of the top 10 enriched pathways. (G and H) UpSet visualization of intersecting sets for genes upregulated (G) or downregulated (H) in APP/PS1 mice with AAV-LEC treatment. Only the differentially expressed genes shared by all 6 cell types are shown in the charts.

**Amino acid sequence of AAV-LEC**:

MDWTWRVFCLLAVAPGAHSDVVMTQSPLSLPVTPGAPASISCRSSQSIVHSNGNTYLEWYLQKPGQSPKLLIYKVSNRFSGVPDRFSGSGSGTDFTLRISRVEAEDVGIYYCFQGSHVPPTFGPGTKLEIKRTSPNSASHSGSAPQTSSAPGSQEVQLVESGGGLVQPGGSLRLSCSASGFTFSSFGMHWVRQAPGKGLEWVAYISSGSSTIYYGDTVKGRFTISRDNAKNSLFLQMSSLRAEDTAVYYCAREGGYYYGRSYYTMDYWGQGTTVTVSSASTKGPSVFPLAPSSKSTSGGTAALGCLVKDYFPEPVTVSWNSGALTSGVHTFPAVLQSSGLYSLSSVVTVPSSSLGTQTYICNVNHKPSNTKVDKKVEPKSCDKTHTCPPCPAPELLGGPSVFLFPPKPKDTLMISRTPEVTCVVVDVSHEDPEVKFNWYVDGVEVHNAKTKPREEQYNSTYRVVSVLTVLHQDWLNGKEYKCKVSNKALPAPIEKTISKAKGQPREPQVYTLPPSRDELTKNQVSLTCLVKGFYPSDIAVEWESNGQPENNYKTTPPVLDSDGSFFLYSKLTVDKSRWQQGNVFSCSVMHEALHNHYTQKSLSLSPGKDYKDDDDK

Signal peptide is shaded, linker is underlined and FLAG tag is red colored.

**DNA sequence of AAV-LEC**:

atggactggacctggagggtgttctgcctgctggccgtggcgcctggtgcgcatagcgacgtggtgatgacccagtcacccctgagcctgccggtgacccctggcgcacctgcttcaatcagctgcagaagctcccaaagcatcgtgcacagcaacggcaacacctacctggaatggtacctgcaaaagccaggccagagcccgaagctgctgatatacaaagtgagcaacaggttcagtggcgtgcccgaccggttctctggcagcggaagcggtacggactttaccctcaggatatctcgcgtggaggcggaggacgtcggaatctattattgcttccaaggttctcacgtcccccccacctttggtcctggaacgaaactggagatcaagaggaccagccccaactccgccagccacagcggctcagctccccagacctctagcgctcctggtagccaagaggtgcagttggtggagagcgggggaggcctggtccagccgggaggctctctgaggctgagctgtagcgcgagcggcttcactttctcatcatttggcatgcactgggtgaggcaggccccaggcaagggcctggagtgggtggcctacatctcaagcggcagctctaccatatactacggcgacaccgtgaagggccgattcaccatcagccgagataacgccaagaacagcctgtttctccagatgagcagtctgagggcagaagacaccgccgtgtactactgtgccagggagggcggctactattacggccggagctactacactatggactactggggacagggcaccacagtgacagtgtctagcgccagcaccaagggtccaagtgtattcccactcgctccaagctctaaatccacgagcggtggtacggcagccctcggatgcctcgtcaaagattattttccagaacctgttacagtatcttggaatagcggcgcgttgaccagtggcgtccatacatttcccgcggtattgcaatcatctggcctctatagtctttcatcagtggtcaccgtcccgagttctagcctgggtacacagacttacatttgcaatgtaaaccataagccaagcaatactaaagttgacaagaaggtcgagcccaaaagctgtgacaagactcatacctgcccgccctgcccagcccccgaacttcttggtggaccgtccgtcttcctctttcctcctaagccgaaggacacgctgatgatctcaaggactcccgaagttacatgtgtggtcgtagatgtctcacacgaggaccccgaggttaaattcaattggtacgtcgacggagtagaagttcataacgccaaaactaagccgagggaggagcaatacaattcaacgtatcgggtcgtaagcgttcttaccgtcctccatcaggattggttgaacggtaaagaatacaagtgcaaagttagcaataaagccctccccgccccgatagaaaagactataagtaaggcgaaaggacaaccaagagaaccccaagtttacactttgcccccctcacgggatgaacttacaaaaaatcaagtctcccttacatgcctggtgaagggtttctacccatcagatatcgcagtggaatgggaatctaacggacaaccagaaaataattacaagacgacgccgccagtcctcgactctgatgggtccttttttttgtactcaaaacttactgtggataaaagtaggtggcaacaaggcaatgtgttctcctgctcagtgatgcatgaggcccttcacaatcattatacccagaaatcattgagcctgagccccggcaaagattacaaggacgacgatgacaagtga
